# Extracellular 2’3’-cGAMP is an immunotransmitter produced by cancer cells and regulated by ENPP1

**DOI:** 10.1101/539312

**Authors:** Jacqueline A. Carozza, Volker Böhnert, Khanh C. Nguyen, Gemini Skariah, Kelsey E. Shaw, Jenifer A. Brown, Marjan Rafat, Rie von Eyben, Edward E. Graves, Jeffrey S. Glenn, Mark Smith, Lingyin Li

## Abstract

2’3’-cyclic GMP-AMP (cGAMP) is characterized as an intracellular second messenger that is synthesized in response to cytosolic dsDNA and activates the innate immune STING pathway. Our previous discovery of its extracellular hydrolase ENPP1 hinted at the existence of extracellular cGAMP. Here, using mass spectrometry, we detected that cGAMP is continuously exported as a soluble factor by an engineered cell line but then efficiently cleared by ENPP1, explaining why it has escaped detection until now. By developing potent, specific, and cell impermeable ENPP1 inhibitors, we detected that cancer cells continuously export cGAMP in culture at steady state and at higher levels when treated with ionizing radiation (IR). In tumors, depletion of extracellular cGAMP using a neutralizing protein decreased tumor-associated immune cell infiltration in a tumor cGAS and host STING dependent manner. Depletion of extracellular cGAMP also abolished the curative effect of IR. Boosting extracellular cGAMP by ENPP1 inhibitors synergizes with IR to shrink tumors in mice. In conclusion, extracellular cGAMP is an anti-cancer immunotransmitter that could be stimulated and harnessed to treat less immunogenic cancers.

## Main

The second messenger 2’3’-cyclic GMP-AMP (cGAMP)^1^ plays pivotal roles in anti-viral and anti-cancer innate immunity. It is synthesized by the enzyme cyclic-GMP-AMP synthase (cGAS)^2^ in response to double-stranded DNA (dsDNA) in the cytosol, which is a danger signal for intracellular pathogens and damaged or cancerous cells^3–7^. cGAMP binds and activates its endoplasmic reticulum (ER) surface receptor Stimulator of Interferon Genes (STING)^8^ to activate production of Type 1 interferons (IFNs). These potent cytokines trigger downstream innate and adaptive immune responses to clear the threat.

In addition to activating STING within its cell of origin, cGAMP can spread to bystander cells through gap junctions in epithelial cells^9^. This cell-cell communication mechanism alerts adjacent cells of the damaged cell and also, unfortunately, accounts for the spreading of drug-induced liver toxicity^10, 11^ and brain metastases^12^. In addition, cytosolic cGAMP can be packaged into budding viral particles and transmitted during the next round of infection^13, 14^. In both transmission modes, cGAMP is not exposed to the extracellular space. Finally, tumor-derived cGAMP has been reported to activate STING in non-cancer cells through unknown mechanisms and eventually activate NK cell responses^15^.

cGAMP is synthesized in the cytosol and cannot passively cross the cell membrane, due to its two negative charges. However, two lines of evidence hinted that cGAMP is exported to the extracellular space to signal other cells. First, we identified a cGAMP hydrolase, ectonucleotide pyrophosphatase phosphodiesterase 1 (ENPP1)^16^, which is the only detectable cGAMP hydrolase reported; however, ENPP1 is annotated as an extracellular enzyme both as a membrane-bound form anchored by a single-pass transmembrane domain and as a cleaved soluble protein in the serum^16^. Second, when added to cell media or injected into tumors, cGAMP and its analogs can cross the cell membrane to activate STING in most cell types^17, 18^; we and others subsequently identified a direct cGAMP importer, SLC19A1.^19, 20^

Here we report direct evidence for cGAMP export by cancers and the role of extracellular cGAMP in anti-cancer immune detection. We subsequently developed nanomolar small molecule inhibitors of ENPP1 and used them to boost extracellular cGAMP concentration, immune infiltration, and tumor progression. Together, we characterize cGAMP as an immunotransmitter that can be harnessed to treat cancer.

## cGAMP is exported from 293T cGAS ENPP1^-/-^ cells as a soluble factor

To test the hypothesis that cGAMP is present extracellularly, we first developed a liquid chromatography-tandem mass spectrometry (LC-MS/MS) method to detect cGAMP from complex mixtures. Using isotopically labeled cGAMP as an internal standard (Extended Data 1a) and optional concentration procedures, we can quantify cGAMP concentrations down to 0.3 nM in both basal cell culture media and serum containing media, and we can quantify intracellular cGAMP concentrations from cell extracts in the same experiment (Extended Data Fig. 1b-d)^21, 22^. We chose to use 293T cells, which express undetectable amounts of cGAS and STING proteins^2, 8, 23^ (Extended Data Fig 1e). By stably expressing mouse cGAS and knocking out *ENPP1* using CRISPR, we created a 293T cGAS ENPP1^low^ cell line and then isolated a single clone to create a 293T cGAS ENPP1^-/-^ cell line. (Extended Data Fig. 1e). We also used serum-free media because serum contains a proteolytically cleaved soluble form of ENPP1^24^. Using this ENPP1-free cell culture system, we detected constant low micromolar basal intracellular cGAMP concentrations in the 293T cGAS ENPP1^-/-^ cells without any stimulation (Fig. 1a), but not in the parent 293T cells (Extended Data Fig. 1f). This was a surprising result when we first observed it in 2016, but can now be explained by multiple subsequent reports showing that cancer cells harbor cytosolic dsDNA as a result of erroneous DNA segregation^25–27^. After replenishing the cells with fresh media, we measured a linear increase of extracellular cGAMP concentrations to 100 nM after 30 h (Fig. 1b). The amount of cGAMP exported is substantial given that after 30 h, the number of cGAMP molecules outside the cells was equal to the number inside (Fig. 1c). We detected negligible amount of cell death based on extracellular lactose dehydrogenate (LDH) activity, suggesting that cGAMP in the media is exported by live cells (Fig. 1d). We calculated the export rate (*v*export) to be 220 molecules cell^-1^ s^-1^ (Fig. 1c). Finally, although human cGAS has been shown to be slower than its mouse counterpart^28^, we measured both intracellular and extracellular cGAMP in human cGAS expressing 293T cells at steady state, which can be further induced with dsDNA transfection (Extended Data Fig. 1g, h).

**Figure 1.**
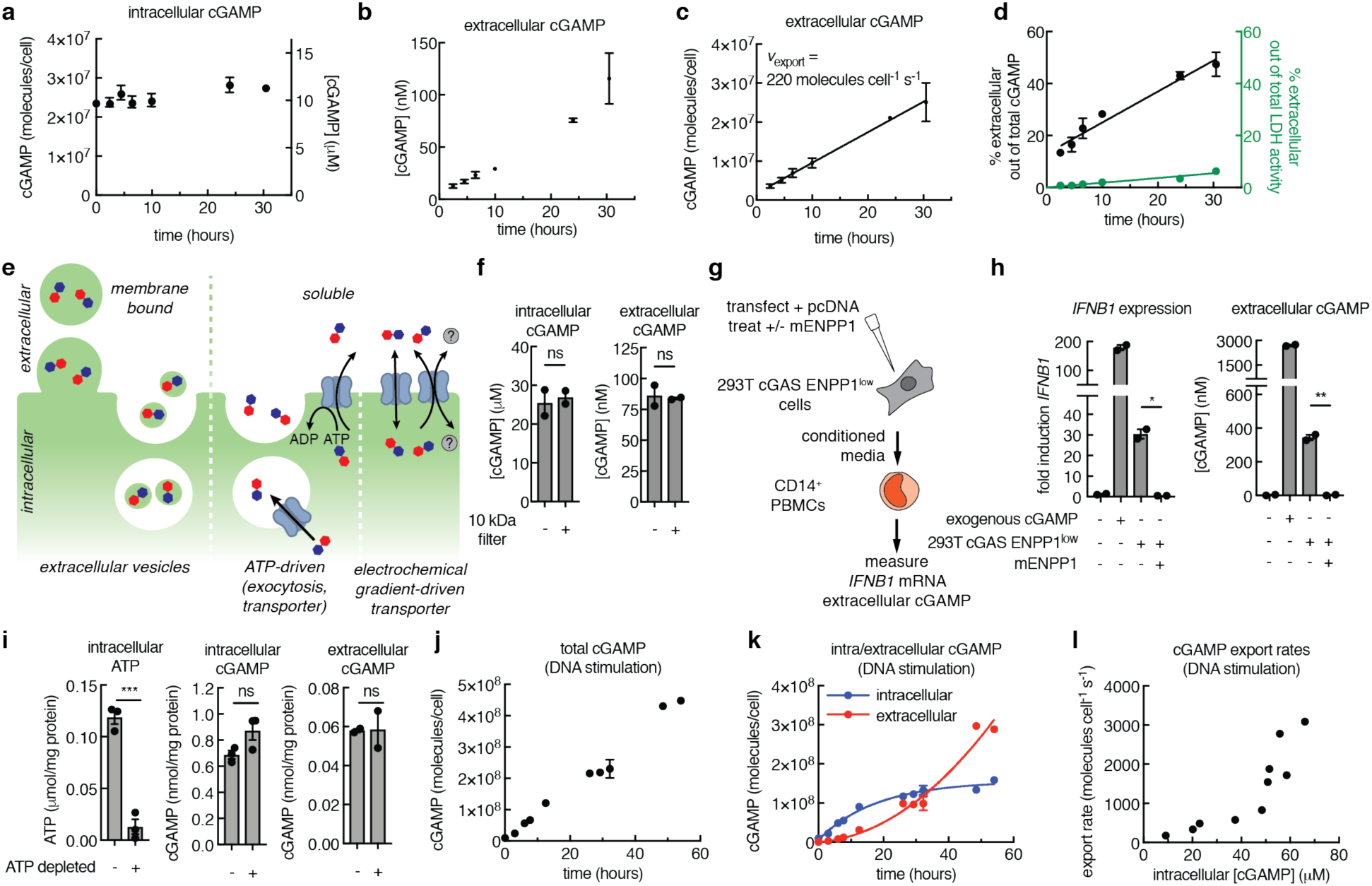
cGAMP is exported from 293T cGAS ENPP1^-/-^ cells as a soluble factor. **a-b**, Intracellular and extracellular concentrations of cGAMP from 293T cGAS ENPP1^-/-^ cells without exogenous stimulation measured using LC-MS/MS. At time 0, cells were replenished with serum-free media. Mean ± SEM (*n* = 2) with some error bars too small to visualize. Data are representative of three independent experiments. **c**, The amount of cGAMP exported per cell over time calculated from data in (**b**). The export rate was calculated using linear regression. **d**, The fraction of extracellular/total cGAMP molecules (left y-axis) calculated from data in (**a**) and (**b**) compared to the fraction of extracellular/total lactate dehydrogenate (LDH) activity as a proxy for cell death (right y-axis). **e**, Potential mechanisms of cGAMP export, including release in extracellular vesicles, exocytosis, ATP-driven transporter, and electrochemical gradient-driven transporter (facilitated diffusion transporter or anti/symporter). **f**, Intracellular and extracellular cGAMP concentrations produced by 293T cGAS ENPP1^-/-^ cells measured before and after passing the media through 10 kDa filters. Mean ± SEM (*n* = 2). Data are representative of two independent experiments. **g**, Schematic of the conditioned media transfer experiment for (**h**). 293T cGAS ENPP1^low^ cells were transfected with 0.5 μg/mL empty pcDNA6 vector and treated ± 20 nM recombinant mouse ENPP1 (mENPP1). Conditioned media from these cells were transferred to primary CD14^+^ human PBMCs. **h**, *IFNB1* mRNA levels were normalized to *CD14* and the fold induction was calculated relative to untreated CD14^+^ cells. Mean ± SEM (*n* = 2). \**P* = 0.04 (one-way ANOVA). cGAMP concentrations were measured in the conditioned media by LC-MS/MS. Mean ± SEM (*n* = 2). \*\**P* = 0.002 (one-way ANOVA). Data are representative of two independent experiments. **i**, 293T cGAS ENPP1^low^ cells were incubated with serum-free ATP depletion media (no glucose, 6 mM 2-deoxy-D-glucose, 5 mM NaN_3_) or serum-free complete media for 1 hour. cGAMP and ATP concentrations were measured by LC-MS/MS. Mean ± SEM (*n* = 2-3). \*\*\**P* < 0.001 (Student’s *t* test). **j-k**, The amount of total (**j**), intracellular (exponential fit), and extracellular (polynomial fit) (**k**) cGAMP over time produced by 293T cGAS ENPP1^-/-^ cells after transfection with empty plasmid pcDNA6 (1.5 μg/mL). Data are representative of two independent experiments. **l**, cGAMP export rates plotted against intracellular concentration of cGAMP after transfection with empty plasmid pcDNA6 (1.5 μg/mL), calculated from data in (**j**) and (**k**).

We have previously shown that there are multiple cGAMP importers and identified SLC19A1 as the first one^19, 20^. We postulate that there are also multiple cGAMP export mechanisms (Fig. 1e). To characterize the cGAMP export mechanism in 293T cells, we first determined whether cGAMP was enclosed in extracellular vesicles (as previously reported^13, 14^) or freely soluble. We filtered conditioned media from 293T cGAS ENPP1^-/-^ cells through a 10 kDa MWCO filter, which retains extracellular vesicles and proteins. cGAMP passed through the filter, suggesting that it is exported as a freely soluble molecule (Fig. 1f). To further confirm that extracellular cGAMP exported by 293T cells is predominantly in a soluble form, we used CD14^+^ human peripheral blood mononuclear cells (PBMCs) as a reporter. These cells have previously been shown to take up soluble cGAMP, which leads to IFN-β production^18^. We observed that CD14^+^ PBMCs respond to submicromolar concentrations of soluble cGAMP by upregulating *IFNB1* (Extended Data Fig. 2a, b). Conditioned media from DNA-transfected cGAS-expressing 293T cGAS ENPP1^low^ cells, but not DNA-transfected 293T cells, induced *IFNB1* expression in CD14^+^ cells, suggesting that the activity is a result of extracellular cGAMP produced by 293T cells (Extended Data Fig. 2c, d). Addition of purified soluble recombinant mouse ENPP1 (mENPP1) (Extended Data Fig. 2e) depleted detectable cGAMP in the conditioned media and also ablated this activity (Fig. 1g, h.) Because soluble ENPP1 (MW = ∼100 kDa) cannot permeate membranes and, thus, can only access soluble extracellular cGAMP, we conclude that 293T cells export soluble cGAMP.

We then determined ATP-dependence of the dominant cGAMP exporter in 293T cells. We depleted ATP for an hour and cell viability was not affected (Extended Data Fig. 2f). In this time period, the intracellular cGAMP concentration remained constant, so did the cGAMP export activity (Fig. 1i). Therefore, in this cell line, cGAMP is not exported by exocytosis or ATP-hydrolyzing pumps, but by an ATP-independent transporter or channel, likely driven by the electrochemical gradient across the cell membrane (Fig. 1e). To further characterize the kinetics of the dominant exporter, we varied intracellular cGAMP concentrations by dsDNA stimulation. Because there is no cGAMP degradation in this ENPP1^-/-^ cell line, the total amount of cGAMP synthesized is the sum of intracellular and extracellular cGAMP (Figure 1j, k). The synthesis rate was linear for the first 12 h and then slowed slightly, possibly due to loss of the dsDNA/cGAS complex over time. By plotting export rates as a function of intracellular cGAMP concentrations, we observed that the *v*export did not plateau in the concentration range we tested (Figure 1l). These kinetics are characteristic of a channel (no measurable *K*m) or an allosterically controlled transporter with a *V*max > 5000 molecules cell^-1^ s^-1^ and a *K*m > 60 μM, as the curve appears sigmoidal instead of hyperbolic (Extended Data Fig. 2g). This characterization will aid in future identification of cGAMP exporter(s).

## Development of a cell impermeable ENPP1 inhibitor to enhance extracellular cGAMP activity

Having established the presence of extracellular cGAMP by carefully removing sources of ENPP1 from culture conditions, we then determined whether intracellular and/or extracellular cGAMP is degraded by ENPP1. Despite its extracellular annotation, it is possible that ENPP1 could flip orientation on the membrane, as reported for a related enzyme CD38^29^, or it could be active when being synthesized in the ER lumen and cGAMP may cross the ER membrane (Fig. 2a). To investigate the localization of ENPP1 activity, we transfected 293T cGAS *ENPP1*^-/-^ cells with human ENPP1 expression plasmid and confirmed its activity in whole cell lysates (Fig. 2b). In intact cells, ENPP1 expression depletes extracellular cGAMP, but does not affect the intracellular cGAMP concentration (Fig. 2c). Therefore, only extracellular cGAMP is regulated by ENPP1 in these cells. However, we cannot exclude the possibility that ENPP1 has intracellular activity in other cell types or under certain stimulations.

**Figure 2.**
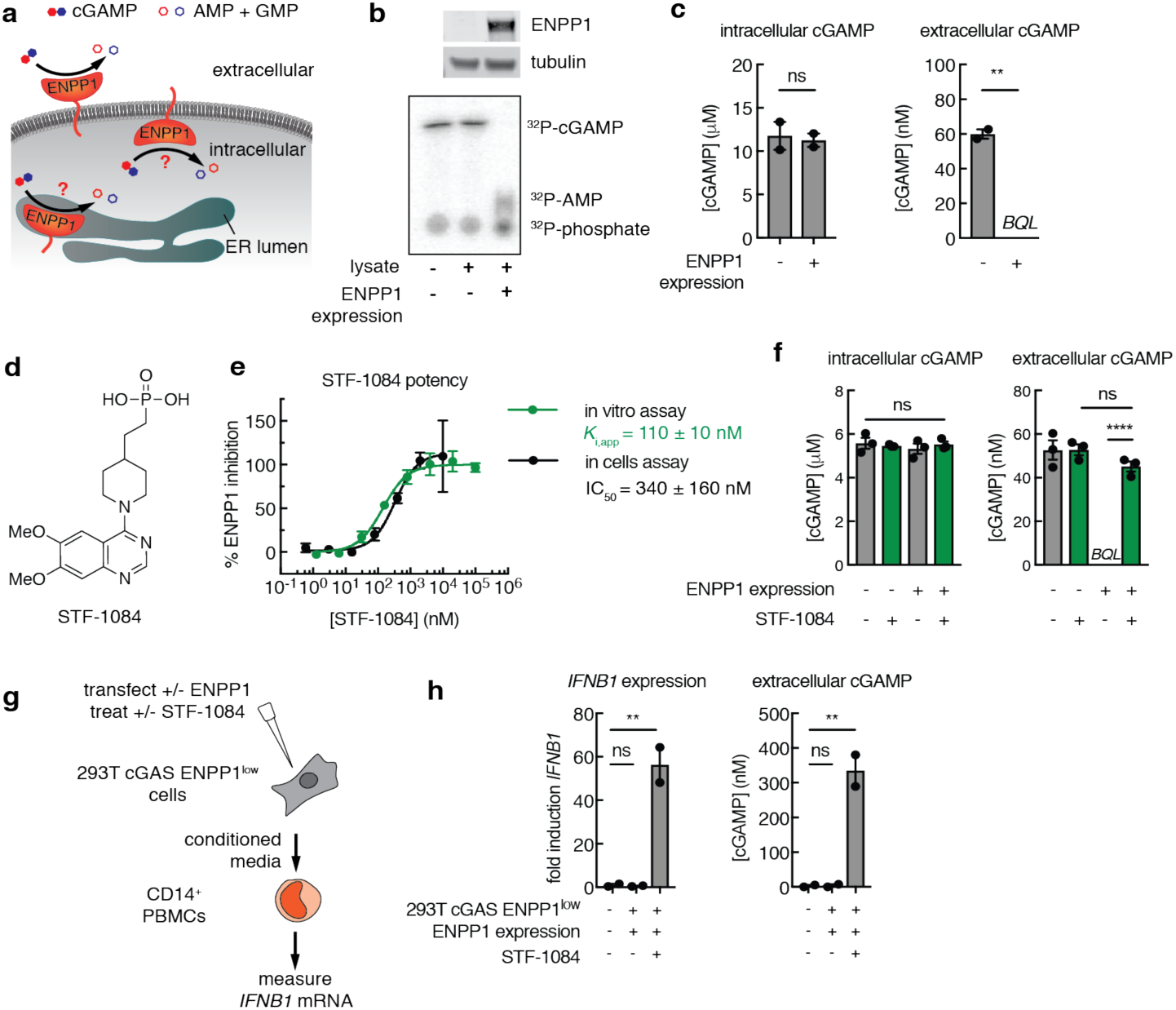
Development of a cell impermeable ENPP1 inhibitor to enhance extracellular cGAMP activity. **a**, Three possible cellular locations of ENPP1 activity. **b**, 293T cGAS ENPP1^-/-^ cells were transfected with empty pcDNA6 vector or vector containing human *ENPP1*. Whole cell lysates were analyzed after 24 h for ENPP1 protein expression using western blot (top) and for ENPP1 ^32^P-cGAMP hydrolysis activity at pH 9.0 using thin layer chromatography (TLC) (bottom). Data are representative of two independent experiments. **c**, Intracellular and extracellular cGAMP concentrations measured using LC-MS/MS. *BQL* = below quantification limit. Mean ± SEM (*n* = 2). \*\**P* = 0.002 (Student’s *t* test). Data are representative of three independent experiments. **d**, Chemical structure of ENPP1 inhibitor STF-1084. **e**, Inhibitory activity of STF-1084 in vitro against purified mouse ENPP1 with ^32^P-cGAMP as the substrate at pH 7.5 (*K*i,app = 110 ± 10 nM) and in cells against human ENPP1 transiently expressed in 293T cGAS ENPP1^-/-^ cells (IC_50_ = 340 ± 160 nM). Extracellular cGAMP levels were analyzed by LC-MS/MS after 24 hours. Mean ± SEM (*n* = 3 independent experiments for in vitro assay, *n* = 2 for in cells assay), with some error bars too small to visualize. **f**, Intracellular and extracellular cGAMP concentrations for 293T cGAS ENPP1^-/-^ cells transfected with empty pcDNA6 vector or vector containing human *ENPP1* in the presence or absence of 10 μM STF-1084 after 24 hours. cGAMP levels were analyzed by LC-MS/MS. *BQL* = below quantification limit. Mean ± SEM (*n* = 3). \*\*\*\**P* < 0.0001 (one-way ANOVA). Data are representative of two independent experiments. **g**, Schematic of the conditioned media experiment for (**h)**. 293T cGAS ENPP1^low^ cells were transfected with vector containing human *ENPP1* and incubated in the presence or absence of STF-1084. Conditioned media from these cells was transferred to primary CD14^+^ human PBMCs. **h**, *IFNB1* mRNA levels were normalized to *CD14* and the fold induction was calculated relative to untreated CD14^+^ cells. Mean ± SEM (*n* = 2). \*\**P* = 0.007 (one-way ANOVA). cGAMP concentrations were measured in the conditioned media by LC-MS/MS. Mean ± SEM (*n* = 2). \*\**P* = 0.006 (one-way ANOVA). Data are representative of two independent experiments.

To study the physiological relevance of extracellular cGAMP, we sought to develop cell impermeable ENPP1 inhibitors that only affect extracellular ENPP1 activity. We first tested a nonspecific ENPP1 inhibitor QS1^30, 31^ (Extended Data Fig. 3a, b). QS1 can inhibit extracellular cGAMP degradation in cells overexpressing ENPP1. However, in ENPP1 null cells, QS1 also increased intracellular cGAMP and decreased extracellular cGAMP concentrations, suggesting that it blocks the cGAMP exporter(s) (Extended Data Fig. 3c). This export blockage activity excludes QS1 as a tool to study extracellular cGAMP. We therefore designed a phosphonate analog, STF-1084, to chelate Zn^2+^ at the ENPP1 catalytic site and to minimize cell permeability and avoid intracellular off-targets (Fig. 2d). STF-1084 is sixty-fold more potent than QS1 (*K*_i,app_ = 110 nM in an in vitro biochemical assay) (Fig. 2e, Extended Data Fig. 3b).

We confirmed that STF-1084 is cell impermeable by performing three independent permeability assays: the parallel artificial membrane permeability assay (PAMPA) (Extended Data Fig. 4a); the intestinal cells Caco-2 permeability assay (Extended Data Fig. 4b); and the epithelial cells MDCK permeability assay (Extended Data Fig. 4c). Compared to control compounds with high cell permeability and low cell permeability, STF-1084 falls into the category of impermeable compounds in all three assays. In addition, it has low activity towards the closely related ectonucleotidases alkaline phosphatase (*K*_i,app_ > 100 μM) and ENPP2 (*K*_i,app_ = 5.5 μM) (Extended Data Fig. 4d). Although we do not expect STF-1084 to have intracellular off-targets due to its low cell permeability, we tested its binding against a panel of 468 kinases to further determine its specificity. Despite its structural similarity to AMP, STF-1084 binds weakly to only two kinases at 1 μM (Extended Data Fig. 4e). STF-1084 also shows high stability (t_1/2_ > 159 min) in both human and mouse liver microsomes, and is non-toxic to primary human PBMCs at 100 μM (Extended Data Fig. 4f). Together, we demonstrated that STF-1084 is a potent, cell impermeable, specific, stable, and non-toxic ENPP1 inhibitor.

Next, we measured the efficacy of STF-1084 in maintaining extracellular cGAMP concentrations of ENPP1 overexpressing 293T cGAS cells and obtained an IC_50_ value of 340 nM, with 10 μM being sufficient to completely block extracellular cGAMP degradation (Fig. 2e). Unlike QS1, STF-1084 had no effect on intracellular cGAMP, demonstrating that it does not affect cGAMP export (Fig. 2f). Finally, we tested the efficacy of STF-1084 in boosting extracellular cGAMP signal detectable to CD14^+^ PBMCs. Conditioned media from ENPP1 overexpressing 293T cGAS cells failed to induce *IFNB1* expression in CD14^+^ cells (Fig. 2g, h). The presence of STF-1084 rescued extracellular cGAMP levels in the media and induction of *IFNB1* expression in CD14^+^ cells (Fig. 2h). STF-1084 had no effect on cytokine production when cGAMP was electroporated into primary human PBMCs (Extended Data Fig. 4g), demonstrating that STF-1084 only boosts extracellular cGAMP signaling by preventing its degradation by ENPP1.

## Cancer cells continuously export cGAMP in culture at steady state and can be induced by ionizing radiation

Chromosomal instability of cancer cells has been reported to lead to micronuclei formation and rupture in the cytosol, and cGAS accumulates at these regions^25–27, 32^. With STF-1084 as a specific extracellular ENPP1 inhibitor, we were poised to test whether cancer cells produce cGAMP and export it. In unstimulated 4T1-luc cells (a mouse triple negative metastatic cancer cell line with a luciferase reporter), we were able to detect 350,000 molecules/cell (∼150 nM) of intracellular cGAMP (Fig. 3a). When we knocked down cGAS in these cells and reduced its protein level by approximately 3-fold (Extended Data Fig. 5a), we also detected a reduction of cGAMP concentration by 3-fold (Fig. 3a), demonstrating that our detection method is indeed measuring cGAMP. In addition, we detected 350,000 molecules/cell extracellular cGAMP in the media after 48 hours (Fig. 3b). Incubating the media with recombinant ENPP1 abolished the signaling, demonstrating that extracellular cGAMP is in a free soluble form (Fig. 3b). Strikingly, when we used STF-1084 to inhibit cell surface and soluble ENPP1 in the cell culture media, we measured 3,000,000 molecules/cell of extracellular cGAMP after 48 hours, which was approximately 10-fold more than intracellular cGAMP, demonstrating that these cells export at least 90% of the cGAMP they synthesize every 48 hours (Fig. 3b). We detected similar levels of extracellular cGAMP in both mouse (4T1-luc, E0771, MC38) and human cancer cells lines (MDA-MB-231 and MCF-7), as well as the immortalized normal mouse mammary gland cells, NMuMG (Fig. 3c). On the contrary, we detected much lower extracellular cGAMP levels in 293T cells, which have very low cGAS expression, and in primary human PBMCs, which have high cGAS expression, but likely no cytosolic dsDNA to stimulate cGAS activity (Fig. 3c). We measured export over time in MC38 cells and it followed linear kinetics as observed in our model 293T cGAS *ENPP1*^-/-^ cells (Fig. 3d). Besides the 4T1-luc cells, the cells we tested (E0771, NMuMG, and Panc02) had intracellular cGAMP levels below our limit of detection, corresponding to less than ∼40 nM intracellular cGAMP (Extended Data Fig. 5b). Interestingly, out of the cell lines that we looked for intracellular cGAMP, only 4T1-luc cells have undetectable amounts of STING protein (Extended Data Fig. 5c). It is possibly that cancer cells upregulate their cGAMP export mechanism(s) to clear intracellular cGAMP to avoid activation of their own STING and subsequent IFN-β production.

**Figure 3.**
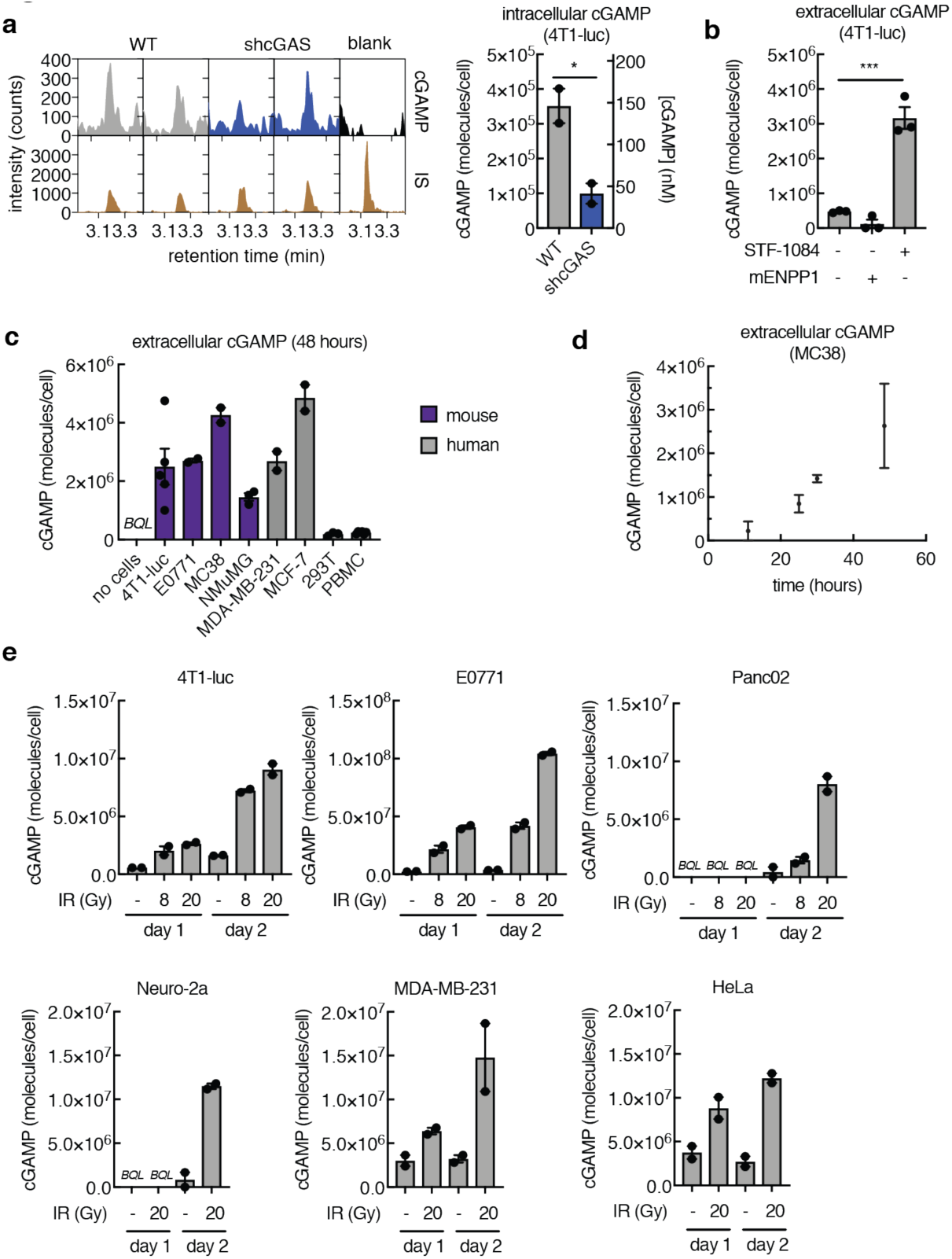
Cancer cells continuously export cGAMP in culture at steady state and can be induced by ionizing radiation (IR). **a**, Intracellular cGAMP in 4T1-luc WT and shcGAS cells without exogenous stimulation measured using LC-MS/MS. Intracellular cGAMP reported in units of molecules/cell and nM (estimated using cell volume = 4 pL). Mean ± SEM (*n* = 2). \**P* < 0.05 (Student’s *t* test). IS = internal standard. **b**, Extracellular cGAMP produced by 4T1-luc cells after 48 hours in serum-containing media in the absence or presence of 50 μM STF-1084. Conditioned media without STF-1084 was also incubated with 10 nM of recombinant mENPP1 overnight. Mean ± SEM (*n* = 3). \*\*\**P* < 0.001 (Student’s *t* test). **c**, Extracellular cGAMP produced by 4T1-luc, E0771, MC38, NMuMG, MDA-MB-231, MCF-7, 293T, and primary human PBMCs measured after 48 h in the presence of 50 μM STF-1084. Mean ± SEM (*n* = 2-6, including data from 2 independent experiments). BQL = below quantification limit. **d**, Extracellular cGAMP produced by MC38 cells over 48 hours. At time 0, cells were refreshed with media supplemented with 50 μM STF-1084. Mean ± SEM (*n* = 2). Data are representative of two independent experiments. **e**, Extracellular cGAMP produced by cancer cell lines 4T1-luc, E0771, Panc02, Neuro-2a, MDA-MB-231, and HeLa after 24 and 48 hours. At time 0, cells were left untreated or treated with IR (8 Gy or 20 Gy) and refreshed with media supplemented with 50 μM STF-1084. Mean ± SEM (*n* = 2). BQL = below quantification limit.

Ionizing radiation (IR) has been shown to increase erroneous chromosomal segregation and cytosolic DNA^25–27, 33, 34^. Indeed, IR increased extracellular cGAMP production in all the cancer cell lines we tested (4T1-luc, E0771, Panc02, Neuro-2a, MDA-MB-231, and HeLa) (Fig. 3e) while causing negligible amounts of cell death (Extended Data Fig. 5d). Interestingly, IR induced more than 10-fold higher extracellular cGAMP in E0771 cells than in other cells lines, despite their similar levels of basal extracellular cGAMP. Together, our data demonstrate that both mouse and human cancer cells, regardless of their tissue of origin, constantly produce and efficiently export cGAMP and can be stimulated with IR to produce more extracellular cGAMP.

## Extracellular cGAMP produced by cancer cells and sensed by host STING is responsible for the curative effect of ionizing radiation

We next directly probed the physiological function of extracellular cGAMP in mouse models. First, to determine the importance of cancer versus host cGAMP, we knocked out *Cgas* in cancer cells (Extended Data Fig. 6a) and utilized *Cgas*^-/-^ and *Sting*^-/-^ mice in the C57BL/6 background (Fig. 4a). We also developed a neutralizing protein agent, which should not be cell permeable, as a tool to specifically sequester extracellular cGAMP. We took advantage of the soluble cytosolic domain of STING (Fig. 4b), which binds cGAMP with a *K*_d_ of 73 ± 14 nM (Fig. 4c). We also generated an R237A^17^ mutant STING as a non-binding STING control (Fig. 4b-d). To test the neutralizing efficacy of these proteins in cell culture, we used CD14^+^ PBMCs. Wild type (WT) STING (neutralizing STING) was able to neutralize extracellular cGAMP with the predicted 2:1 stoichiometry, while the non-binding STING had no effect even at a 200-fold higher concentration (Fig. 4e). We further validated the use of the STING proteins as tools to probe extracellular cGAMP function in primary mouse bone marrow cells, where dosing with neutralizing STING ablated cytokine production compared to non-binding STING (Extended Data Fig. 6b)

**Figure 4.**
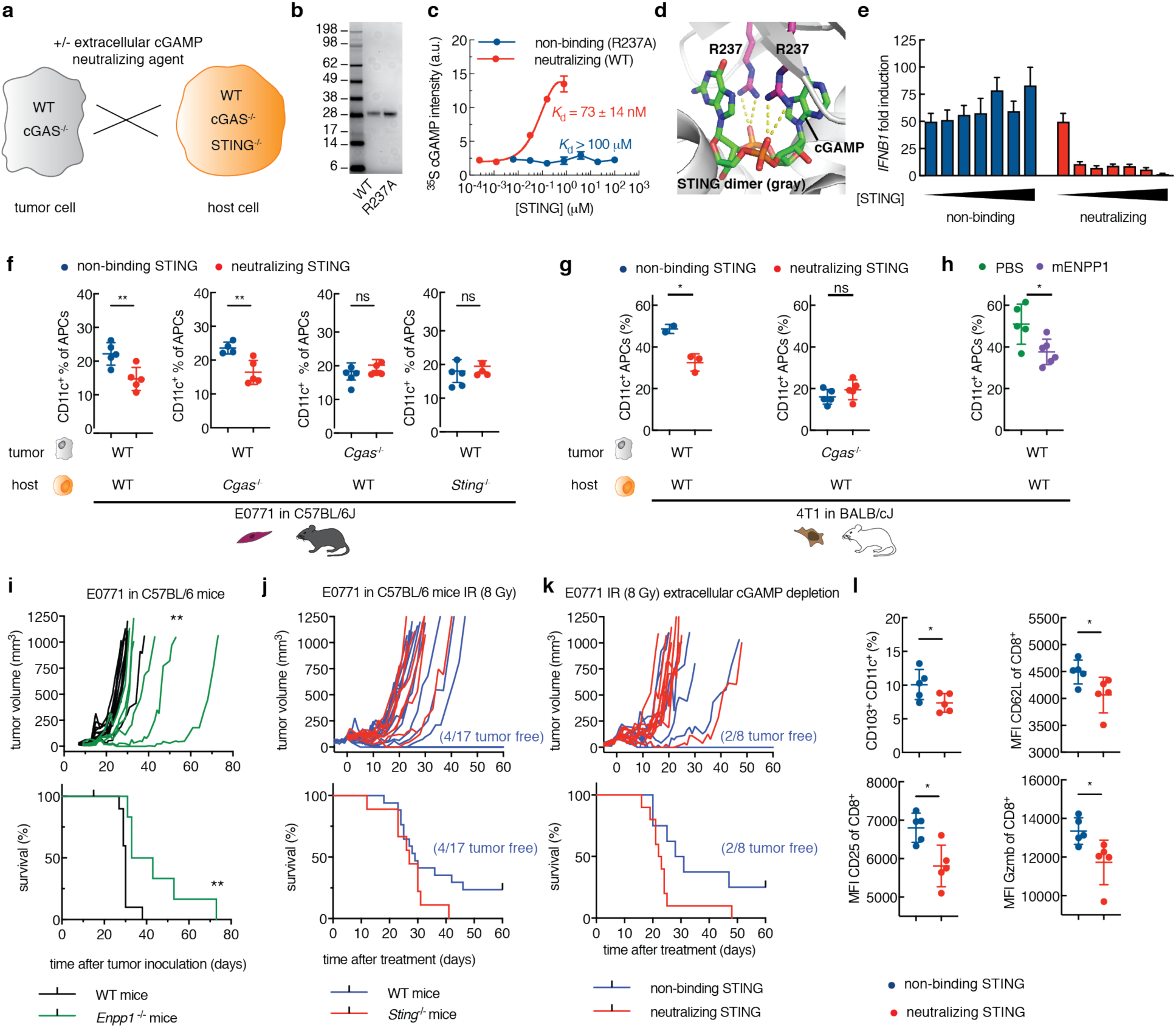
Extracellular cGAMP produced by cancer cells and sensed by host STING is responsible for the tumor shrinkage effect of ionizing radiation. **a**, Experimental setup to assess the role of extracellular cGAMP in vivo. **b**, Coomassie gel of the cytosolic domain of recombinant mouse WT and R237A STING. **c**, Binding curves for the cytosolic domain of mouse WT STING (neutralizing) and R237A STING (non-binding) determined by a membrane binding assay using radiolabeled ^35^S-cGAMP as the probe. Mean ± SEM (*n* = 2 from two independent experiments). **d**, Crystal structure of mouse WT STING in complex with cGAMP with R237 highlighted in pink (PDB ID 4LOJ). **e**, *IFNB1* mRNA fold induction in CD14^+^ PBMCs treated with 2 μM cGAMP in the presence of neutralizing or non-binding STING (2 μM to 100 μM, 2.5-fold dilutions). Mean ± SEM (*n* = 2 technical qPCR replicates). **f**, WT or *Cgas*^-/-^ E0771 cells (1×10^6^) were orthotopically injected into WT, *Cgas*^-/-^, or *Sting*^-/-^ C57BL/6J mice on day 0. Neutralizing STING or non-binding STING was intratumorally injected on day 2. Tumors were harvested and analyzed by FACS on day 3. Samples were gated on cells in FSC-A/SSC-A, singlets (FSC-W), living cells, CD45^+^, MHC II^+^ (APCs), CD11c^+^ populations. (Non-binding STING: WT mice *n* = 5; *Cgas*^-/-^ mice *n* = 4; *Cgas*^-/-^ cells *n* = 5; *Sting*^-/-^ mice *n* = 5. Neutralizing STING: WT mice *n* = 5; *Cgas*^-/-^ mice *n* = 5; *Cgas*^-/-^ cells *n* = 5; *Sting*^-/-^ mice *n* = 4). Mean ± SD. \*\**P* < 0.01. (Welch’s *t* test). **g**, WT or *Cgas*^-/-^ 4T1-luc cells (1×10^6^) were orthotopically injected into WT BALB/cJ mice, and the same procedure was performed as in (**f**). Percent CD11c^+^ cells of total MHC II^+^ (APCs). (Non-binding STING: WT cells *n* = 2; *Cgas*^-/-^ cells *n* = 5; neutralizing STING: WT cells *n* = 3; *Cgas*^-/-^ cells *n* = 5). \**P* < 0.05. (Welch’s *t* test). **h**, 4T1-luc cells (1×10^6^) were orthotopically injected into WT BALB/cJ mice on day 0. PBS (*n* = 5) or recombinant mouse ENPP1 (mENPP1) (*n* = 6) was intratumorally injected on day 2. Tumors were harvested and analyzed by FACS on day 3. Mean ± SD. \**P* < 0.05. (Welch’s *t* test). **i**, E0771 cells (5×10^4^) were orthotopically injected into WT (*n* = 10) or Enpp1^-/-^ (*n* = 6) C57BL/6J mice. Tumor volume and survival were monitored. \*\**P* < 0.01. *P* value for tumor volume determined by pairwise comparisons using post hoc tests with a Tukey adjustment and for Kaplan Meier curve determined using the Log-rank Mantel-Cox test. **j**, E0771 cells (5×10^4^) were orthotopically injected into WT (*n* = 10) or *Sting^-^*^/-^ (*n* = 9) C57BL/6J mice. The tumors were treated with IR (8 Gy) when they reached 100 ± 20 mm^3^. Tumor volume and survival were monitored without further treatment. **k**, E0771 cells (5×10^4^) were orthotopically injected into WT C57BL/6J mice. The tumors were treated with IR (8 Gy) when they reached 100 ± 20 mm^3^ and injected with non-binding (*n* = 8) or neutralizing STING (*n* = 10) every other day for the duration of the experiment. Tumor volume and survival were monitored. **l**, E0771 cells (5×10^4^) were orthotopically injected into WT C57BL/6J mice. The tumors were treated with IR (8 Gy) when they reached 100 ± 20 mm^3^ and injected with non-binding (*n* = 5) or neutralizing STING (*n* = 5) on days 2 and 4 after IR. Tumors were harvested and analyzed by FACS on day 5. Mean ± SD. \**P* = < 0.05. (Welch’s *t* test).

We established E0771 orthotopic tumors in mice, followed by intratumoral injection of neutralizing STING to deplete extracellular cGAMP, and excision of the tumors to stain for tumor-associated leukocytes. In WT E0771 tumors, compared to non-binding STING, neutralizing STING significantly decreased the CD11c^+^ (dendritic cell or DC) population and the CD103^+^ CD11c^+^ (conventional type I dendritic cell or cDC1) population^35^ in the CD45^+^ MHC II^+^ (tumor-associated antigen presenting cell or APC) population, suggesting that extracellular cGAMP can be detected by the immune system (Fig. 4f and Extended Data Fig. 6c, d). Extracellular cGAMP depletion also diminished the CD11c^+^ population when tumors are grown in *Cgas*^-/-^ mice, suggesting that host cells do not contribute significantly to extracellular cGAMP production (Fig. 4f). In contrast, extracellular cGAMP depletion did not affect the CD11c^+^ or CD103^+^ CD11c^+^ population when *Cgas*^-/-^ E0771 cells (multiple clones were pooled to achieve clean knockout but minimize clonal effects) or *Sting*^-/-^ mice were used (Fig. 4f and Extended Data Fig. 6d). Depleting extracellular cGAMP did not affect the F4/80^+^ (macrophage) population in the APC population in any of the experiments (Extended Data Fig. 6e). Together, our data demonstrate that cancer cells, but not host cells, are the dominant producers of extracellular cGAMP, which is then sensed by host STING and leads to infiltration of DCs, in particular cross-presenting cDC1s. We also tested the orthotopic 4T1-luc tumor model in the BALB/c background. Although *Cgas* and *Sting* knockout strains have not been established in this background, we knocked out *Cgas* in the 4T1-luc cells and pooled multiple clones. Intratumoral injection of neutralizing STING into the WT 4T1-luc tumors significantly decreased the tumor-associated CD11c^+^ population in the CD45^+^ MHC II^+^ population (Fig. 4g). In contrast, extracellular cGAMP depletion had no effect in *Cgas*^-/-^ 4T1-luc tumors. As an orthogonal approach, we depleted extracellular cGAMP by intratumoral injection of mENPP1 protein (Extended Data Fig. 2e) and again observed diminished CD11c^+^ cells in the CD45^+^ MHC II^+^ population (Fig. 4h). Together, our results demonstrate that extracellular cGAMP is produced by cancer cGAS and sensed by host STING, which leads to increased immune cell infiltration. We next tested whether this basal level of extracellular cGAMP can also lead to a tumor shrinkage effect. Long term administration of neutralizing STING compared to the non-binding STING control did not alter the course of tumor progression in the E0771 model (Extended Data Fig. 6f), suggesting that endogenous ENPP1-mediated degradation was sufficient to abolish the anti-cancer effect of cancer-derived extracellular cGAMP. E0771 cells do not have particularly high ENPP1 activity (Extended Data Fig. 7a), but ENPP1 is also expressed on host cells and present in the serum as a soluble form^16, 24, 36^. We therefore implanted WT E0771 cells into the *Enpp1*^-/-^ mice and, indeed, observed slower tumor growth, suggesting that host ENPP1 promotes tumor growth in this model. (Fig. 4i).

We next tested the physiological role of endogenous extracellular cGAMP when stimulated by IR, without ENPP1 inhibition. It was previously reported that IR exerts tumor-shrinkage effects in a host STING-dependent manner^37^ and activates cGAS-dependent IFN-β production in cancer cells^25, 26, 38^. Indeed, IR treatment induced cytokine production in both 4T1-luc and E0771 cells (Extended Data Fig. 7b). Since E0771 cells export high levels of cGAMP upon IR treatment (Fig. 3e), we investigated the role of extracellular cGAMP in the tumor shrinkage effect of IR in the E0771 breast tumor model. We did not observe a significant increase of dead cells or cleaved caspases in both CD45^-^ and CD45^+^ cells in response to treatment with IR (8 Gy) at 24h, suggesting that IR does not directly kill cancer or immune cells (Extended Data Fig. 7c). Treating established E0771 tumors (100 ± 20 mm^3^) with 8 Gy of IR resulted in tumor-free survival in 4 out of 17 mice, demonstrating the efficacy of IR in this model (Fig. 4j). No tumor-free survival was observed in *Sting*^-/-^ mice, confirming that the curative effect of IR depends on host STING in this model, but probably not cytokines produced by cancer cells (Fig. 4j). We sequestered extracellular cGAMP by injecting neutralizing STING for the duration of the experiment with non-binding STING as a control. Remarkably, depletion of extracellular cGAMP completely abolished the curative effect of IR (Fig. 4k), suggesting that extracellular cGAMP-induced host STING activation accounts for the curative effect of IR in this model.

To probe the cellular mechanism of extracellular cGAMP, we excised established tumors five days after treating them with IR (8 Gy) followed by extracellular cGAMP depletion and analyzed the immune cell populations (Extended Data Fig. 7d). The amount of tumor infiltrating CD11c^+^ and F4/80^+^ cells were not significantly altered when depleting extracellular cGAMP with neutralizing STING versus the non-binding STING control (Extended Data Fig. 7e). However, the CD103^+^ CD11c^+^ subpopulation decreased, indicating weakened ability in cross-presentation when extracellular cGAMP was depleted (Fig. 4l). Although the numbers of CD8^+^ T cells were not altered significantly (Extended Data Fig. 7e), expression levels of the activation markers CD62L, CD25, and Granzyme B were dampened when extracellular cGAMP was depleted (Fig. 4l). Together, our results suggest that IR cures tumors by increasing the extracellular cGAMP level, which then directly or indirectly increases tumor infiltrating cross-presenting cDC1s, leading to activation of CD8^+^ cytotoxic T cells.

## ENPP1 inhibitors synergize with IR to shrink tumors

ENPP1 is highly expressed in some breast cancers and its level has been correlated with poor prognosis^39–41^. High ENPP1 expression may be a mechanism that breast cancers utilize to deplete extracellular cGAMP and dampen immune detection. We measured ENPP1 activities in three triple negative breast cancer cells 4T1-luc, E0771, and MDA-MB-231, with MDA-MB-231 and 4T1-luc exhibiting high ENPP1 activities (Extended Data Fig. 7a). We, therefore, chose the 4T1-luc mouse model to probe the effect of ENPP1 on tumor immune detection, growth, and responses to treatment. We first tested the effect of ENPP1 on tumor infiltrating dendritic cells. We knocked out ENPP1 in 4T1-luc cells, validated the clones by their lack of enzymatic activity (commercially available ENPP1 antibodies are not sensitive enough to validate true knockout), and pooled multiple clones to minimize clonal effects (Extended Data Fig. 8a). Three days after orthotopic 4T1-luc tumor implantation, we excised the tumors and analyzed their tumor-associated leukocyte compositions. *Enpp1*^-/-^ tumors have a larger tumor-associated CD11c^+^ population than WT tumors when left untreated or treated with IR (20 Gy) (Fig. 5a). To further test that this effect was due to increased extracellular cGAMP, but not potential membrane scaffolding effects of ENPP1, any unidentified intracellular activity, or because these 4T1-luc cells express Cas9 due to the CRISPR knock out procedure, we used our cell impermeable ENPP1 inhibitor. We intratumorally injected STF-1084 immediately after IR treatment and tested its effect after 24h. Indeed, compared to vehicle control, STF-1084 mirrored the effect of *Enpp1* knockout on increasing the tumor-associated CD11c^+^ population (Fig. 5b).

**Figure 5.**
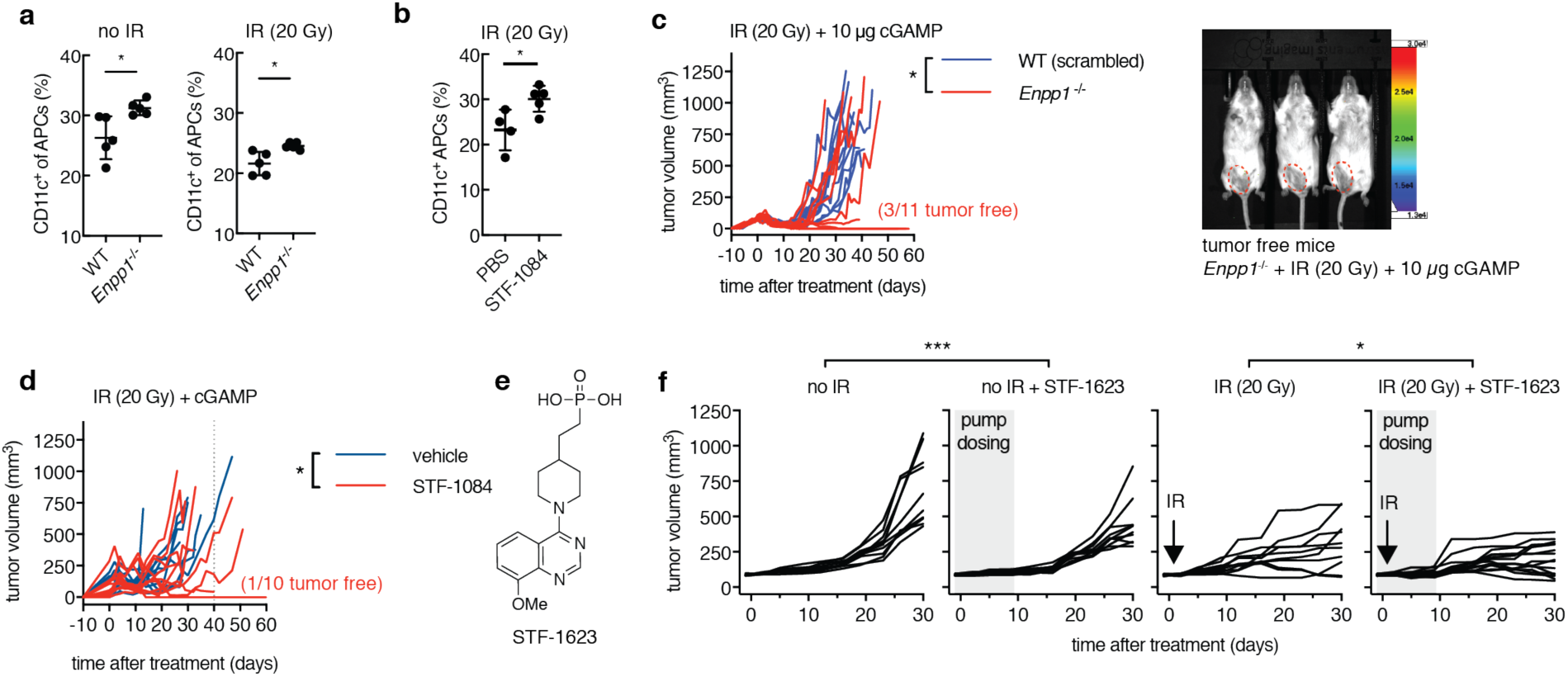
ENPP1 inhibitors synergize with IR to shrink tumors. **a**, 4T1-luc WT or *Enpp1*^-/-^ cells (1×10^6^) were orthotopically injected into WT BALB/cJ mice on day 0. Tumors were left untreated (*n* = 5) or treated with IR (20 Gy) (*n* = 5) on day 2. Tumors were harvested and analyzed by FACS on day 3. \**P* < 0.05. (Welch’s *t* test). **b**, 4T1-luc cells (1×10^6^) were orthotopically injected into WT BALB/cJ mice on day 0. Tumors were treated with 20 Gy IR and intratumorally injected with PBS (*n* = 4) or STF-1084 (*n* = 5) on day 2. Tumors were harvested and analyzed by FACS on day 3. \**P* < 0.05 (Welch’s *t* test). **c**, Established 4T1-luc WT (harboring scrambled sgRNA) or *Enpp1*^-/-^ cells (100 ± 20 mm^3^) were treated once with IR (20 Gy) followed by three intratumoral injections of 10 µg cGAMP on day 2, 4, and 7 after IR (*n* = 10 for scrambled 4T1-luc, *n* = 11 for Enpp1^-/-^ 4T1-luc). Tumor volumes are shown. \**P* < 0.05. *P* value determined by pairwise comparisons using post hoc tests with a Tukey adjustment at day 20. In the *Enpp1*^-/-^ 4T1-luc + IR (20 Gy) + 10 µg cGAMP treatment group, 3/11 (27%) mice are tumor free verified by bioluminescent imaging (tumor area outline in red). **d**, Established 4T1-Luc tumors (100 ± 20 mm^3^) were treated once with IR (20 Gy) followed by three intratumoral injections of 10 µg cGAMP alone or 10 µg cGAMP + 100 µL of 1 mM STF-1084 on day 2, 4, and 7 after IR (*n* = 9 per treatment group). Tumor volumes are shown. \**P* < 0.05. *P* value determined by pairwise comparisons using post hoc tests with a Tukey adjustment at day 40. **e**, Chemical structure of ENPP1 inhibitor STF-1623. **f**, Mice bearing established subcutaneous Panc02 tumors (100 ± 20 mm^3^) were implanted with a subcutaneous pump containing STF-1623 (50 mg/kg/day) on day 0 and left untreated or treated with IR (20 Gy) on day 1. No IR: *n* = 10, no IR + STF-1623: *n* = 10, IR (20 Gy): *n* = 10, IR (20 Gy) + STF-1623: *n* = 15. Pumps were removed on day 8. Tumor volumes are shown. *P* value determined by pairwise comparisons using post hoc tests with a Tukey adjustment. \**P* < 0.05. \*\*\**P* < 0.001.

We then tested the effect of ENPP1 expressed by 4T1-luc on tumor rejection. We did not observe significant growth delay of 4T1-luc *Enpp1*^-/-^ tumors compared to WT tumors harboring a scrambled sgRNA and Cas9 when they were left untreated, or treated with IR or intratumoral cGAMP injections individually (Extended Data Fig. 8b-d) Strikingly, a 30% cure rate was achieved when we treated *Enpp1*^-/-^ tumors with a combination of IR and cGAMP injections, whereas no mice inoculated with WT tumors were cured (Fig. 5c). This demonstrates that ENPP1 expressed on the surface of 4T1 cells was sufficient to abolish the tumor shrinkage effect of combination therapy. Together, we have demonstrated that ENPP1 expressed on both cancer cells (using the 4T1 model) and by the host (using the E0771 model) play a role in clearing extracellular cGAMP and promote tumor growth.

Although it is still an open question whether cancer or host ENPP1 plays a bigger role, small molecule inhibitors should, in principle, inhibit both. We tested the ENPP1 inhibitor STF-1084 in this combination therapy. STF-1084 has fast pharmacokinetics. Without optimizing its route of administration and pharmacokinetic properties, we intratumorally injected it into orthotopic 4T1-luc tumors when they reached 100 mm^3^. STF-1084 synergized with IR and cGAMP to significantly delay tumor progression, and resulted in tumor free survival of 1/10 mice. (Fig. 5d).

Other than breast cancers, pancreatic cancers also express high levels of ENPP1 (Extended Data Fig. 8e). To be able to access these tumors that are not easily accessible through intratumoral injections, we sought to develop an analog of STF-1084 that can be administered systemically, thus also inhibiting both host and cancer cell ENPP1. We developed STF-1623 (Fig. 5e), with improved *K*_i_ of 16 ± 7 nM (Extended data Fig 9a). We performed a similar suite of assays on STF-1623 as we did on STF-1084, confirming that it affected only extracellular cGAMP concentrations (Extended data Fig 9b) and bore a similar safety and stability profile (Extended Data Fig. 9c-f). Importantly, STF-1623 shows an improved pharmacokinetic profile compared to STF-1084 for systemic dosing. With subcutaneous injections, we can achieve >100 nM plasma concentrations after 24 hours (Extended Data Fig. 9g).

We then tested STF-1623 in the Panc02, syngeneic, subcutaneous pancreatic tumor model. STF-1623 delayed tumor progression as a single agent and synergized with IR to achieve tumor regression in 33% of mice (Fig. 5f). Together, our results demonstrate that the anti-tumor effect of extracellular cGAMP can be enhanced by inhibiting its degradation enzyme ENPP1. STF-1623 serves as a starting point for new classes of anti-cancer agents that can synergize with the endogenous extracellular cGAMP exported by cancer cells and induced by IR.

## Extracellular cGAMP in cancer

Here we report direct evidence of cGAMP export. Cell–cell cGAMP transfer through gap junctions and viral particles has been demonstrated previously. We provide the first in vitro and in vivo evidence that cGAMP can travel through the extracellular space (Extended Data Fig. 10). In all the cell types we have tested, cGAMP can be exported at various levels, suggesting that cGAMP export is not a cancer specific phenomenon and cGAMP exporters are likely expressed in most cell types. Since chromosomal instability and aberrant cytosolic dsDNA are cancer-intrinsic properties^42, 43^ and cancer cells rarely inactivate cGAS^27^, we reason that cGAMP overproduction and increased export may also be properties intrinsic to cancer cells. Since no cytosolic cGAMP hydrolase has been identified and ENPP1 cannot degrade intracellular cGAMP, export is currently the only known mechanism by which cGAMP is removed from the cytosol, and represents another way to turn off intracellular STING signaling in addition to ubiquitin-mediated STING degradation^44^. This clearance mechanism, however, exposes cancer cells to immune detection.

Indeed, our results demonstrate that cGAMP exported by cancer cells is a danger signal detected by the immune system. It is well known that neoantigens from cancer cells are presented by APCs to cross prime cytotoxic CD8^+^ T cells that eventually perform cancer-specific killing^7, 18^. However, it is less understood how APCs initially detect cancer cells. It has been shown that immunogenic tumors release dsDNA as a danger signal to CD11c^+^ dendritic cells^7, 45^. The evidence for IFN as a danger signal is mixed; one study showed that cancer cells respond to their own cytosolic dsDNA induced by IR and produce IFNs as a danger signal^46^, whereas other studies have reported that cancer cells can lose their ability to make IFN via the STING pathway, or even repurpose the STING pathway to aid in metastasis^27, 47^. A recent study showed that the catalytic activity of cancer cGAS correlates with anti-cancer immunity in the B16 melanoma model in a host STING dependent manner^15^, suggesting that cGAMP could be transferred from cancer cells to host cells, with unknown mechanism. Here, we provide direct evidence that cancer cells produce soluble extracellular cGAMP as a danger signal, which leads to increased numbers of dendritic cells, specifically cross-presenting type I conventional dendritic cells, and cytotoxic T cell activation in the tumor microenvironment. cGAMP export is an important mode of cGAMP communication among cells that are not physically connected but are in close proximity. Unlike cytokines, cGAMP cannot travel long distance in the extracellular space without being degraded and/or diluted to below its effective concentrations. This property is shared with neurotransmitters and qualifies cGAMP as the first identified immunotransmitter.

## Acknowledgments

This work is dedicated to Dr. T. Mitchison to celebrate his 60^th^ birthday and his remarkable achievements in understanding biochemical mechanisms of the cell. He taught Dr. L. Li the power of definitive experiments. We thank the following institutions and people for their help: Stanford Small Animal Imaging Facility, S. Ergun for ^35^S cGAMP, C. Patel for specificity assays, F. Sunden for enzyme assays, R.Stabler for chemical synthesis, N. Weng for protein purification, Dr. C. Walsh, Dr. D. Herschlag, and all members of the Li lab for helpful discussions. Flow cytometry analysis for this project was done on instruments in the Stanford Shared FACS Facility. Data was collected on an instru
ment in the Shared FACS Facility obtained using NIH S10 Shared Instrument Grant S10RR027431-01.

## Author Contributions

J.A.C., V.B., K.E.S., M.S., and L.L. designed the study. J.A.C., V.B., K.E.S., K.N., G.S., J.A.B., M.R., and L.L. performed experiments and analyzed data. R.E. performed statistical analysis. J.A.C., V.B., and L.L. wrote the manuscript. J.S.G. supervised K.N.; E.E.G. supervised M.R and R.E. All authors discussed the findings and commented on the manuscript.

## Data and materials availability

All data are available in the main text or supplementary information.

## Materials and Methods

### Reagents and antibodies

[*α*-^32^P]ATP (800 Ci/mmol, 10 mCi/mL, 250 μCi) and [^35^S]ATP*α*S (1250 Ci/mmol, 12.5 mCi/mL, 250 μCi) were purchased from Perkin Elmer. Adenosine triphosphate, guanosine triphosphate, adenosine-^13^C_10_,^15^N_5_, 5’-triphosphate, 4-nitrophenyl phosphate, and bis(4-nitrophenyl) phosphate were purchased from Sigma-Aldrich and are >98% atomically pure. 2’3’-cGAMP and isotope labeled cGAMP were synthesized as described previously^19^. Caco-2 assay was purchased from Cyprotex. Kinome screens were conducted by Eurofins (data visualized using TREE*spot*™ Software Tool and reprinted with permission from KINOME*scan*®, a division of DiscoveRx Corporation. ©DiscoveRX Corporation 2010).

. PAMPA and MDCK permeability assays were conducted by Quintara Discovery. Total protein content was quantified using the BCA assay (Thermo Fisher). Cell viability was quantified using the CellTiterGlo assay (Promega) or the lactate dehydrogenase assay (Pierce, Thermo Fisher). Mouse CXCL10 production was quantified with the mouse CXCL10/IP-10/CRG-2 DuoSet ELISA (R&D Systems) and the TMB substrate reagent set (BD Bioscience). Full length human *ENPP1* was cloned into pcDNA6 vector. QS1 was synthesized as previously described^31^. The following monoclonal antibodies were used for western blotting: rabbit anti-cGAS (D1D3G Cell Signaling, 1:1,000) rabbit anti-mouse cGAS (D2O8O Cell Signaling, 1:1,000), mouse anti-tubulin (DM1A Cell Signaling, 1:2,000), and rabbit anti-STING (D2P2F Cell Signaling, 1:1,000), IRDye 800CW goat anti-rabbit (LI-COR, 1:15,000), and IRDye 680RD goat anti-mouse (LI-COR, 1:15,000).

### Mammalian cell lines and primary cells

293T, NMuMG, MDA-MB-231, HeLa, MCF-7, and Neuro-2a were procured from ATCC, Panc02 was procured from the DTP/DCTD/NCI Tumor Repository, E0771 was procured from CH3 BioSystems, 4T1-luciferase (4T1-luc) was a gift from Dr. Christopher Contag^48^, and HEK293S GnT1^-^ expressing secreted mENPP1 was a gift from Dr. Osamu Nureki^49^. All cell lines were maintained in a 5% CO_2_ incubator at 37°C. 293T, Neuro-2a, MDA-MB-231, and HeLa were maintained in DMEM (Corning Cellgro) supplemented with 10% FBS (Atlanta Biologics) (v/v) and 100 U/mL penicillin/streptomycin (P/S) (ThermoFisher). NMuMG and MCF-7 were maintained in DMEM supplemented with 10% FBS, 100 U/mL P/S, and 10 μg/mL bovine insulin (Millipore Sigma). MC38 cells were maintained in DMEM supplemented with 10% FBS, and 100 U/mL P/S, 2 mM Glutamine, 0.1 mM nonessential amino acids, 1 mM sodium pyruvate, and 10mM HEPES (all Gibco). 4T1-luc, and Panc02 were maintained in RPMI (Corning Cellgro) supplemented with 10% FBS, 100 U/mL P/S. E0771 cells were maintained in RPMI supplemented with 10% FBS, 100U/ml P/S, and 10 mM HEPES. Primary human peripheral blood mononuclear cells (PBMCs) (Stanford Blood Center) were isolated by subjecting enriched buffy coat from whole blood to a Percoll density gradient. CD14^+^ PBMCs were isolated using CD14^+^ MicroBeads (Miltenyi). PBMCs were cultured in RPMI supplemented with 2% human serum and 100 U/mL P/S. Primary mouse bone marrow cells were isolated by opening the ends of the femur and tibia removing cells by centrifugation^50^. Bone marrow cells were culture in RPMI supplemented with 10% FBS and 100 U/mL P/S.

#### Making cell lines

293T cells were virally transfected to stably express mouse or human cGAS. 293T cGAS ENPP1^low^ cells were created by viral transfection of CRISPR sgRNA targeting human *ENPP1* (5’-CACCGCTGGTTCTATGCACGTCTCC-3’), and 293T mcGAS ENPP1^-/-^ cells were selected after single cell cloning from this pool. 4T1 and E0771 *Cgas*^-/-^ cells were created by viral transfection of CRISPR sgRNA (using lentiCRISPRv2-blast, Addgene plasmid #83480^51^) targeting mouse *Cgas* (5’-CACCGGAAGGGGCGCGCGCTCCACC-3’). Cells were single cell cloned, and multiple clean knockouts were pooled after verification by western blotting. 4T1-luc *Enpp1*^-/-^ and scrambled cells were created by viral transfection of CRISPR sgRNAs (using lentiCRISPRv2-blast) targeting mouse *Enpp1* (5’-GCTCGCGCCCATGGACCT-3’ and 5’-ATATGACTGTACCCTACGGG -3’) or a scrambled sequence. Cells were selected with 0.5 – 2 μg/mL blasticidin, single cell cloned, and multiple clean knockouts were pooled after verification by activity assay. Alternatively, 4T1-luc *Enpp1*^-/-^ and scrambled cells were created by transient transfection with Lipofectamine 3000 of the same CRISPR sgRNAs as above or a scrambled sequence (using PX458, Addgene plasmid # 48138), followed by single cell cloning of GFP positive cells. Multiple clean knockouts were pooled after verification by activity assay 4T1-luc shcGAS cells were created by viral transfection of shRNA (5’-CAGGATTGAGCTACAAGAATAT-3’)^46^ using the plasmid pGH188. Cells harboring the shRNA were selected with 0.5 – 2 μg/mL blasticidin, sorted for GFP expression, and were used as a pool for experiments.

### Expression and purification of recombinant proteins

sscGAS was produced as described previously^19^. mENPP1 was produced as described previously^49, 52^.

Mouse STING (residues 139-378) was inserted into the pTB146 His-SUMO vector (a generous gift from T. Bernhard, Harvard Medical School) and expressed in Rosetta cells. Cells were grown in 2xYT medium with 100 μg/mL ampicillin and induced when the OD_600_ reached 1 with 0.75 mM IPTG at 16 °C overnight. All subsequent procedures using proteins and cell lysates were performed at 4 °C. Cells were pelleted and lysed in 50 mM Tris pH 7.5, 400 mM NaCl, 10 mM imidazole, 2 mM DTT, and protease inhibitors (cOmplete, EDTA-free protease inhibitor cocktail Roche). Cells were lysed by sonication and the lysate was cleared by ultracentrifugation at 50,000 rcf for 1 hour. The cleared supernatant was incubated with HisPur cobalt resin (ThermoFisher Scientific; 1 mL resin per 1 L bacterial culture) for 30 minutes. The resin-bound protein was washed with 50 column volumes of 50 mM Tris pH 7.5, 150 mM NaCl, 2% triton X-114, 50 CV of 50 mM Tris pH 7.5, 1 M NaCl (each wash was set to a drip rate of 1 drop/2-3 seconds and took 2-3 hours), and 20 column volumes of 50 mM Tris pH 7.5, 150 mM NaCl. Protein was eluted from resin with 600 mM imidazole in 50 mM Tris pH 7.5, 150 mM NaCl. Fractions containing His-SUMO-STING were pooled, concentrated, and dialyzed against 50 mM Tris pH 7.5, 150 mM NaCl while incubating with the SUMOlase enzyme His-ULP1 to remove the His-SUMO tag overnight. The solution was incubated with the HisPur cobalt resin again to remove the His-SUMO tag, and STING was collected from the flowthrough. Protein was dialyzed against 20 mM Tris pH 7.5, loaded onto a HitrapQ anion exchange column (GE Healthcare) using an Äkta FPLC (GE Healthcare), and eluted with a NaCl gradient. Fractions containing STING were pooled and buffer exchanged into PBS and snap frozen at -80 °C until use.

### Liquid chromatography-tandem mass spectrometry

Measurement of cGAMP: Cyclic GMP-^13^C_10_,^15^N_5_-AMP was used as an internal standard at 0.5-1 μM. Samples were analyzed for cGAMP, ATP, and GTP content on a Shimadzu HPLC (San Francisco, CA) with an autosampler set at 4°C and connected to an AB Sciex 4000 QTRAP (Foster City, CA). A volume of 10 μL was injected onto a Biobasic AX LC column, 5 μm, 50 × 3 mm (Thermo Scientific). The mobile phase consisted of 100 mM ammonium carbonate (A) and 0.1% formic acid in acetonitrile (B). Initial condition was 90% B, maintained for 0.5 min. The mobile phase was ramped to 30% A from 0.5 min to 2.0 min, maintained at 30% A from 2.0 min to 3.5 min, ramped to 90% B from 3.5 min to 3.6 min, and maintained at 90% B from 3.6 min to 5 min. The flow rate was set to 0.6 mL/min. The mass spectrometer was operated in electrode spray positive ion mode with the source temperature set at 500°C. Declustering and collision-induced dissociation were achieved using nitrogen gas. Declustering potential and collision energy were optimized by direct infusion of standards. For each molecule, the MRM transition(s) (*m/z*), DP (V), and CE (V) are as follows: ATP (508 > 136, 341, 55), GTP (524 > 152, 236, 43), cGAMP (675 > 136, 121, 97; 675 > 312, 121, 59; 675 > 152, 121, 73), internal standard cyclic GMP-^13^C_10_,^15^N_5_-AMP (690 > 146, 111, 101; 690 > 152, 111, 45; 690 > 327, 111, 47).

Measurement of STF-1084 and STF-1623: Measurements were performed using a Q-Exactive FT-mass spectrometer (Thermo) equipped with Vanquish uHPLC. Samples were diluted in water with 0.1% formic acid and injected onto a Phenomenex Synergi Hydro-RP column (4 μm particle size 2 mm ID, 30 mm length). The column compartment was at ambient temperature. The flow rate was 0.5 mL/min. Mobile phase A was water with 0.1% formic acid; mobile phase B was acetonitrile with 0.1% formic acid. Each run was five minutes; the gradient was as follows: 0-0.5 min 0% B, 0.5 to 2 min linear from 0 to 95% B, 2 to 3.5 min hold at 95% B, 3.5 to 3.6 min from 95% B to 0% B and 3.6 to 5 min at 0% B to re-equilibrate the column. The first minute of the run was diverted to waste and minutes 1 through 4.8 were sent to the mass spectrometer for analysis. Detection on the Q-Exactive was performed in positive mode between 100–1000 m/z, using an acquisition target of 1E6 with maximum IT of 100 ms at a resolution of 70,000. Quantification was done using TraceFinder 4.1 software (Thermo).

### Export assay in 293T cGAS ENPP1^-/-^ cells

293T cGAS ENPP1^-/-^ cells were plated in tissue culture treated plates coated with PurCol (Advanced BioMatrix). At the start of the experiment, the media was gently removed and replaced with serum-free DMEM supplemented with 1% insulin-transferrin-selenium-sodium pyruvate (ThermoFisher) and 100 U/mL penicillin-streptomycin. At indicated times, the media was removed and the cells were washed off the plate with cold PBS. Both the media and cells were centrifuged at 1000 rcf for 10 minutes at 4 °C. The cells were lysed in 30 to 100 μL of 50:50 acetonitrile:water supplemented with 500 nM internal standard, and centrifuged at 15,000 rcf for 20 minutes at 4 °C to remove the insoluble fraction. If no concentration was necessary, an aliquot of media was removed, supplemented with internal standard at 500 nM and 20% formic acid. If concentration/extraction was necessary for detecting < 4 nM cGAMP, the media was acidified with 0.5% acetic acid and supplemented with internal standard (the appropriate amount for a final concentration of 1 μM in 100 μL). Media was applied to HyperSep Aminopropyl SPE columns (ThermoFisher Scientific) to enrich for cGAMP as described previously^22^. Eluents were evaporated to dryness and reconstituted in 50:50 acetonitrile:water. The media and cell extract were submitted for mass spectrometry quantification of cGAMP, ATP, and GTP.

### Transfection stimulation of 293T cGAS ENPP1^-/-^ cells

293T cGAS ENPP1^-/-^ cells were transfected with Fugene 6 (Promega) according to manufacturer’s instructions plus indicated concentrations of pcDNA6 plasmid DNA (empty or containing human *ENPP1*). 24 hours following transfection, the export assay was conducted as described above.

### ATP depletion experiment in 293T cGAS ENPP1^-/-^ cells

293T cGAS ENPP1^-/-^ cells were incubated with serum-free ATP depletion media (no glucose, 6 mM 2-deoxy-D-glucose, 5 mM NaN_3_) or serum-free complete media for 1 hour. The export assay processing was conducted as described above.

### Conditioned media transfer

293T cGAS ENPP1^low^ cells were plated and transfected with plasmid DNA as described above. 24 hours following transfection, media was changed to RPMI + 2% human serum + 1% penicillin-streptomycin, ± 2 μM cGAMP, ± 20 nM recombinant mENPP1, or ± 50 uM STF-1084. 24 hours following media change, the conditioned media was removed from the 293T cGAS ENPP1^low^ cells and incubated with freshly isolated CD14^+^ PBMCs. Gene expression of CD14^+^ PBMCs was analyzed 14-16 h later.

### PBMC electroporation of cGAMP and treatment with ENPP1 inhibitor

PBMCs (2 × 10^6^) were resuspended in electroporation buffer (90 mM Na_2_HPO_4_, 90 mM NaH_2_PO_4_, 5 mM KCl, 10 mM MgCl_2_, 10 mM sodium succinate) with or without 200 nM cGAMP. Cells were electroporated in a cuvette with a 0.2 cm electrode gap (Bio-Rad) and electroporated using program U-013 on a Nucleofector II device (Lonza) and immediately transferred to fresh media with or without ENPP1 inhibitor.

### RT-PCR analysis

Total RNA was extracted using Trizol (Thermo Fisher Scientific) and reverse transcribed with Maxima H Minus Reverse Transcriptase (Thermo Fisher Scientific). Real-time RT-PCR was performed in duplicate with AccuPower 2X Greenstar qPCR Master Mix (Bioneer) on a 7900HT Fast Real-Time PCR System (Applied Biosystems). Data were normalized to *CD14*, *ACTB*, or *GAPDH* expression (human) and *Actb* expression (mouse) for each sample. Fold induction was calculated using ΔΔCt. Primers for human *IFNB1*: fwd (5’-AAACTCATGAGCAGTCTGCA-3’), rev (5’-AGGAGATCTTCAGTTTCGGAGG-3’); human *CXCL10*^9^: fwd (5’-TCTGAATCCAGAATCGAAGG-3’), rev (5’-CTCTGTGTGGTCCATCCTTG-3’) human *CD14*: fwd (5’-GCCTTCCGTGTCCCCACTGC-3’), rev (5’-TGAGGGGGCCCTCGACG-3’); human *ACTB*: fwd (5’-GGCATCCTCACCCTGAAGTA-3’), rev (5’-AGAGGCGTACAGGGATAGCA-3’); human *GAPDH*: fwd (5’-CCAAGGTCATCCATGACAAC-3’); rev (5’-CAGTGAGCTTCCCGTTCAG-3’); mouse *Cxcl10*^9^: fwd (5’-CTCTGTGTGGTCCATCCTTG-3’), rev (5’-GTGGCAATGATCTCAACACG-3’), mouse *Actb*^9^: fwd (5’-AGCCATGTACGTAGCCATCC-3’), rev (5’-CTCTCAGCTGTGGTGGTGAA-3’)

### ^32^P-cGAMP degradation TLC assay

Radiolabeled ^32^P cGAMP was synthesized by incubating unlabeled ATP (1 mM) and GTP (1 mM) doped with ^32^P-ATP with 2 μM purified recombinant porcine cGAS in 20 mM Tris pH 7.5, 2 mM MgCl_2_, 100 μg/mL herring testes DNA) overnight at room temperature, and the remaining nucleotide starting materials were degraded with alkaline phosphatase for 4 h at 37 °C. Cell lysates were generated by scraping and lysing 1×10^6^ cells (293T) or 10×10^6^ cells (4T1-luc, E0771, and MDA-MB-231) in 100μL of 10 mM Tris, 150 mM NaCl, 1.5 mM MgCl_2_, 1% NP-40, pH 9.0. For 4T1-luc, E0771, and MDA-MB-231, total protein concentration of lysate was measured using the BCA assay (Pierce, Thermo Fisher), and samples were normalized so the same amount of protein was used for each lysate reaction. The probe ^32^P-cGAMP (5 μM) was incubated with mENPP1 (20 nM) or whole cell lysates in 100 mM Tris, 150 mM NaCl, 2 mM CaCl_2_, 200 μM ZnCl_2_, pH 7.5 or pH 9.0 for the indicated amount of time. To generate inhibition curves, 5-fold dilutions of ENPP1 inhibitor was included in the reaction. Degradation was evaluated by TLC as previously described^16^. Plates were exposed on a phosphor screen (Molecular Dynamics) and imaged on a Typhoon 9400 and the ^32^P signal was quantified using ImageJ. Inhibition curves were fit to obtain IC_50_ values with Graphpad Prism 7.03. IC_50_ values were converted to K_i,app_ values using the Cheng-Prusoff equation K_i,app_ = IC_50_/(1+[S]/K_m_).

### ALPL and ENPP2 inhibition assays

Inhibition assays for other ectonucleotidases were performed by incubating reaction components in 96-well plate format at room temperature and monitoring production of 4-nitrophenolate by measuring absorbance at 400 nM in a platereader (Tecan). ALPL: 0.1 nM ALPL, 2 μM 4-nitrophenyl phosphate, and various concentrations of inhibitor in buffer pH 9.0 containing 50 mM Tris, 20 μM ZnCl_2_, 1 mM MgCl_2_ at room temperature. ENPP2: 2 nM ENPP2, 500 μM bis(4-nitrophenyl) phosphate, and various concentrations of inhibitor in buffer pH 9.0 containing 100 mM Tris, 150 mM NaCl, 200 μM ZnCl_2_, 2 mM CaCl_2_.

### Intracellular and extracellular cGAMP measurement in cancer cell lines

Cells were refreshed at time 0 with media supplemented with 50 μM STF-1084 (for IR experiments, cancer cell lines were also exposed to 8 Gy or 20 Gy of *γ*-radiation using a cesium source. At indicated times, media was collected and centrifuged at 1,000 rcf; cells were trypsinized off the plate, quenched with serum-containing media, washed with PBS, and counted. Cells were then lysed with 80% water + 20% methanol + 2% acetic acid and centrifuged at 15,000 rcf. cGAMP was enriched from the media and cell supernatant as described above using the HyperSep Aminopropyl SPE columns and submitted for mass spectrometry quantification.

### Mouse models (4T1-luc, E0771, Panc02)

Seven- to nine-week-old female BALB/c mice (Jackson Laboratories) were inoculated with 5 × 10^4^ or 5 × 10^5^ 4T1-luc or 4T1-luc ENPP1^-/-^ cells suspended in 50 μL of PBS into the fifth mammary fat pad. Seven- to nine-week-old female C57B6/J WT,STING^gt/gt^ (referred to as *Sting*^-/-^), or *Enpp1*^-/-^ mice (Jackson Laboratories) were inoculated with 5 × 10^4^ E0771 cells suspended in 50 μL of PBS into the fifth mammary fat pad. Five- to nine-week-old female C57B6/J mice were inoculated with 3 × 10^6^ Panc02 cells suspended in 100 μL of PBS subcutaneously into the right hind flank. When tumor volume (determined by length^2^ × width / 2) reached 100 ± 20 mm^3^, tumors were irradiated with 20 Gy (4T1-luc and Panc02 models) or 8Gy (E0771 model) using a 225 kVp cabinet X-ray irradiator filtered with 0.5 mm Cu (IC-250, Kimtron Inc., CT). Anaesthetized animals were shielded with a 3.2 mm lead shield with a 15 × 20 mm aperture where the tumor was placed. For Panc02, the mice were implanted subcutaneously between the scapulae with an osmotic pump (Alzet 1002) containing a solution of 200 mg/mL STF-1623 in PBS or PBS alone one day prior to IR. Pumps were removed 8 days after implantation. Treatments after irradiation were administered as specified. Tumor volumes were recorded and analyzed in a generalized estimation equation in order to account for the within mouse correlation. Pair-wise comparisons of the treatment groups at each time point were done using post hoc tests with a Tukey adjustment for multiple comparisons. Animal death was plotted in a Kaplan Meier curve using Graphpad Prism 7.03 and statistical significance was assessed using the Log-rank Mantel-Cox test. Mice were maintained at Stanford University in compliance with the Stanford University Institutional Animal Care and Use Committee regulations and procedures were approved by the Stanford University administrate panel on laboratory animal care, or mice were maintained by Crown Biosciences in accordance with their regulations on animal laboratory care.

### FACS analysis of tumors

Seven- to nine-week-old female BALB/c WT (4T1-luc tumors) or C57BL/6 (E0771 tumors) WT, *Cgas*^-/-^, or *Sting*^gt/gt^ (referred to as *Sting*^-/-^) mice (Jackson Laboratories) were inoculated with 1 × 10^6^ tumor cells suspended in 50 μL of PBS into the fifth mammary fat pad. Two days after injection, tumors were intratumorally injected with 100 μL of 1 mM STF-1084 in PBS or with PBS alone. For experiments using STING and mENPP1, 100 μL of 100 μM neutralizing STING or non-binding STING (R237A) or 700 nM mENPP1 or PBS were injected intratumorally.

Alternatively, seven- to nine-week-old female C57BL/6 (Jackson Laboratories) were inoculated with 5 × 10^4^ E0771 tumor cells suspended in 50 μL of PBS into the fifth mammary fat pad. After reaching 100 ± 20 mm^3^, tumors were irradiated with 8 Gy. Two and four days after IR, tumors were intratumorally injected with 100 μL of 100 μM neutralizing STING or non-binding STING (R237A).

On the next day, the tumor was extracted and incubated in RPMI + 10% FBS with 20 μg/mL DNase I type IV (Sigma-Aldrich) and 1 mg/mL Collagenase from Clostridium histolyticum (Sigma-Aldrich) at 37 °C for 30 min. Tumors were passed through a 100 μm cell strainer (Sigma-Aldrich) and red blood cells were lysed using red blood cell lysis buffer (155 mM NH_4_Cl, 12 mM NaHCO_3_, 0.1 mM EDTA) for 5 min at room temperature. Cells were stained with Live/Dead fixable near-IR dead cell staining kit (Thermo Fisher Scientific), Fc-blocked for 10 min using TruStain fcX and subsequently antibody-stained with CD8α, CD11c, CD45, CD62L, F4/80, Granzyme B, I-A/I-E (all Biolegend), CD3ε, CD25 (both eBioscience), and CD103 (BD Biosciences).

Caspase activity was detected after red blood cell lysis using the FAM-FLICA Poly Caspase Assay Kit (ImmunoChemistry Technologies) according to the manufacturer’s description.

Cells were analyzed using an SH800S cell sorter (Sony), an LSR II (BD Biosciences), or an Aurora (Cytek). Data was analyzed using FlowJo V10 software (Treestar) and Prism 7.04 software (Graphpad) for statistical analysis and statistical significance was assessed using the unpaired t test with Welch’s correction.

### In vivo imaging

Mice were injected ip with 3mg XenoLight D-luciferin (Perkin-Elmer) in 200 µl water and imaged using a Lago X in vivo imaging system (Spectral Instruments Imaging). Object height was set to 1.5cm, binning to 4, FStop to 1.2, and the exposure time was 120s. Images were analyzed using aura 2.0.1 software (Spectral Instruments Imaging).

### Synthesis of STF-1084

#### Preparation of dimethyl (E)-(2-(1-benzylpiperidin-4-yl)vinyl)phosphonate

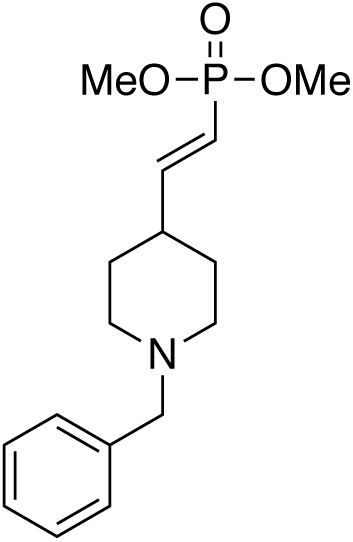

Sodium hydride (2.16 g, 54.11 mmol) was carefully added to a stirred solution of bis(dimethoxyphosphoryl)methane (11.42 g, 49.19 mmol) in toluene (100 mL) at room temperature. The reaction mixture was then placed under an atmosphere of nitrogen and a solution of 1-benzylpiperidine-4-carbaldehyde (10 g, 49.19 mmol) in toluene (50 mL) was slowly added keeping the temperature below 40 °C. The resulting mixture was left to stir at room temperature for 16 h and then quenched by the addition of aqueous saturated ammonium chloride solution. The organic phase was separated, washed with brine, dried (MgSO_4_) and evaporated to dryness. Chromatography (120 g SiO_2_; 5 to 100% gradient of EtOAc in hexanes) provided dimethyl (*E*)-(2-(1-benzylpiperidin-4-yl)vinyl)phosphonate (6.2 g, 16%) as a colorless oil.

LC-MS: m/z = 309.8 [M+H]^+^

^1^H NMR (500 MHz, Chloroform-*d*) – δ 7.47–7.21 (m, 5H), 6.86–6.73 (m, 1H), 5.65–5.53 (m, 1H), 3.75 (s, 3H), 3.70 (s, 3H), 3.52 (s, 2H), 2.98–2.87 (m, 2H), 2.22–2.11 (m, 1H), 2.09–1.99 (m, 2H), 1.79–1.70 (m, 2H) and 1.54–1.44 (m, 2H).

#### Preparation of dimethyl (2-(piperidin-4-yl)ethyl)phosphonate

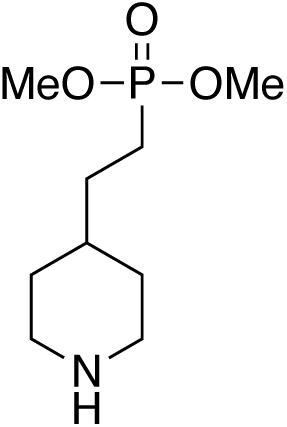

To a mixture of dimethyl (*E*)-(2-(1-benzylpiperidin-4-yl)vinyl)phosphonate (6.2 g, 20.0 mmol) in ethanol (80 mL) was added Pd(OH)_2_/C (0.5 g). The mixture was exchanged with hydrogen gas for three times and stirred under hydrogen balloon at room temperature for 12 h. The mixture was filtered through Celite® and evaporated to dryness to give 4.4 g (100% yield in ∼90% purity) of dimethyl (2-(piperidin-4-yl)ethyl)phosphonate as colorless oil.

^1^H NMR (500 MHz, Chloroform-*d*) δ 3.75 (s, 3H), 3.73 (s, 3H), 3.08 (dt, *J* = 12.5, 3.2 Hz, 2H), 2.58 (td, *J* = 12.2, 2.5 Hz, 2H), 1.80–1.60 (m, 4H), 1.60–1.50 (m, 2H), 1.38 (m, 1H), 1.11 (qd, *J* = 12.1, 4.0 Hz, 2H).

#### Preparation of dimethyl (2-(1-(6,7-dimethoxyquinazolin-4-yl)piperidin-4-yl)ethyl)phosphonate

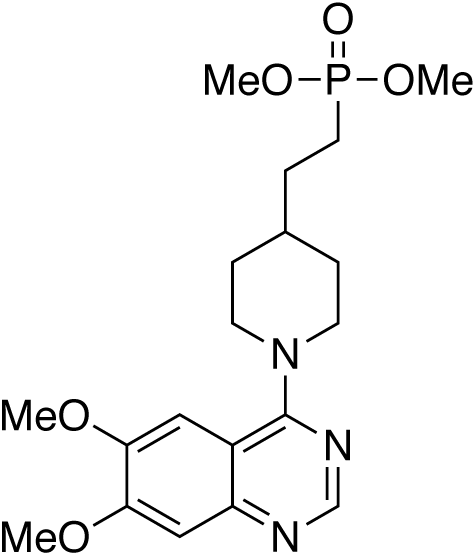

Diisopropylethylamine (0.6 g, 8.9 mmol) was added to a mixture of dimethyl (2-(piperidin-4-yl)ethyl)phosphonate (1.1 g, 4.9 mmol) and 4-chloro-6,7-dimethoxyquinazoline (1.0 g, 4.5 mmol) in isopropyl alcohol (20 mL). After stirring at 90 °C for 3 h, the reaction mixture was cooled and evaporated to dryness. Purification of silica gel (5% MeOH in dichloromethane) provided dimethyl (2-(1-(6,7-dimethoxyquinazolin-4-yl)piperidin-4-yl)ethyl)phosphonate (755 mg, 37%) as oil.

LC-MS: m/z = 410.25 [M+H]^+^

^1^H NMR (500 MHz, CDCl_3_) δ 8.65 (s, 1H), 7.23 (s, 1H), 7.09 (s, 1H), 4.19 (dq, *J* = 14.0, 2.9, 2.4 Hz, 2H), 4.02 (s, 3H), 3.99 (s, 3H), 3.77 (s, 3H), 3.75 (s, 3H), 3.05 (td, *J* = 12.8, 2.3 Hz, 2H), 1.93 – 1.77 (m, 4H), 1.67 (ddd, *J* = 14.1, 9.5, 5.9 Hz, 3H), 1.46 (qd, *J* = 12.2, 3.7 Hz, 2H).

#### Preparation of dimethyl (2-(1-(6,7-dimethoxyquinazolin-4-yl)piperidin-4-yl)ethyl)phosphonic acid hydrogen bromide salt (STF-1084)

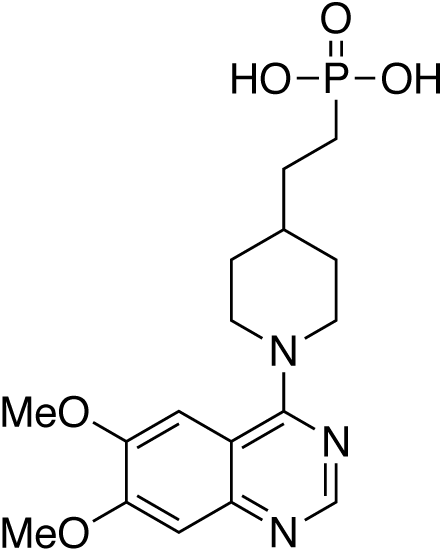

Bromotrimethylsilane (3.67 g, 24 mmol) was added to a cooled solution of dimethyl (2-(1-(6,7-dimethoxyquinazolin-4-yl)piperidin-4-yl)ethyl)phosphonate (3.25 g, 7.94 mmol) in chloroform (60 mL) that was cooled by an ice bath. The reaction mixture was allowed to warm to room temperature and after 90 minutes was quenched by the addition of methanol (20 mL). The mixture was evaporated to dryness under reduced pressure and then solvated in methanol (100 mL). The reaction mixture was concentrated to half volume, filtered to remove precipitate, and then evaporated to dryness. The residue was crystalized with dichloromethane, filtered and dried under vacuum to give dimethyl (2-(1-(6,7-dimethoxyquinazolin-4-yl)piperidin-4-yl)ethyl)phosphonic acid (2.1 g, 69%).

LC-MS: m/z = 381.8 [M+H]^+^

^1^H NMR (500 MHz, DMSO-*d*_6_) δ 8.77 (s, 1H), 7.34 (s, 1H), 7.23 (s, 1H), 4.71 (d, *J* = 13.1 Hz, 2H), 3.99 (s, 3H), 3.97 (s, 3H), 3.48 (t, *J* = 12.7 Hz, 2H), 3.18 (s, 1H), 1.97–1.90 (m, 2H), 1.62–1.43 (m, 4H), 1.40–1.27 (m, 2H).

### Synthesis of STF-1623

(2-(1-(8-methoxyquinazolin-4-yl)piperidin-4-yl)ethyl)phosphonic acid (STF-1623) was prepared according to the same synthetic procedure as STF-1084, using the 4-chloro-8-methoxyquinazoline instead of the 4-chloro-6,7-dimethoxyquinazoline in the preparation of dimethyl (2-(1-(8-dimethoxyquinazolin-4-yl)piperidin-4-yl)ethyl)phosphonate.

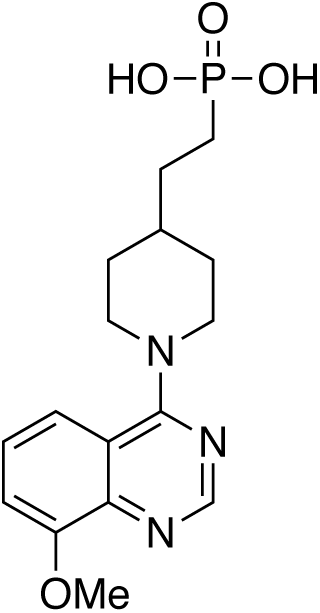

LC-MS: m/z = 352.1 [M+H]^+^

^1^H NMR (400 MHz, DMSO-*d*_6_): δ 8.62 (s, 1H), 7.63–7.53 (m, 3H), 6.65 (d, J = 12.4 Hz, 2H), 4.02 (s, 3H), 3.39–3.36 (m, 2H), 1.92 (m, 2H), 1.58-1.56 (m, 1H), 1.54–1.45 (m, 4H) and 1.35–1.30 (m, 2H).

## Extended Data Figures

**Extended Data Figure 1.**
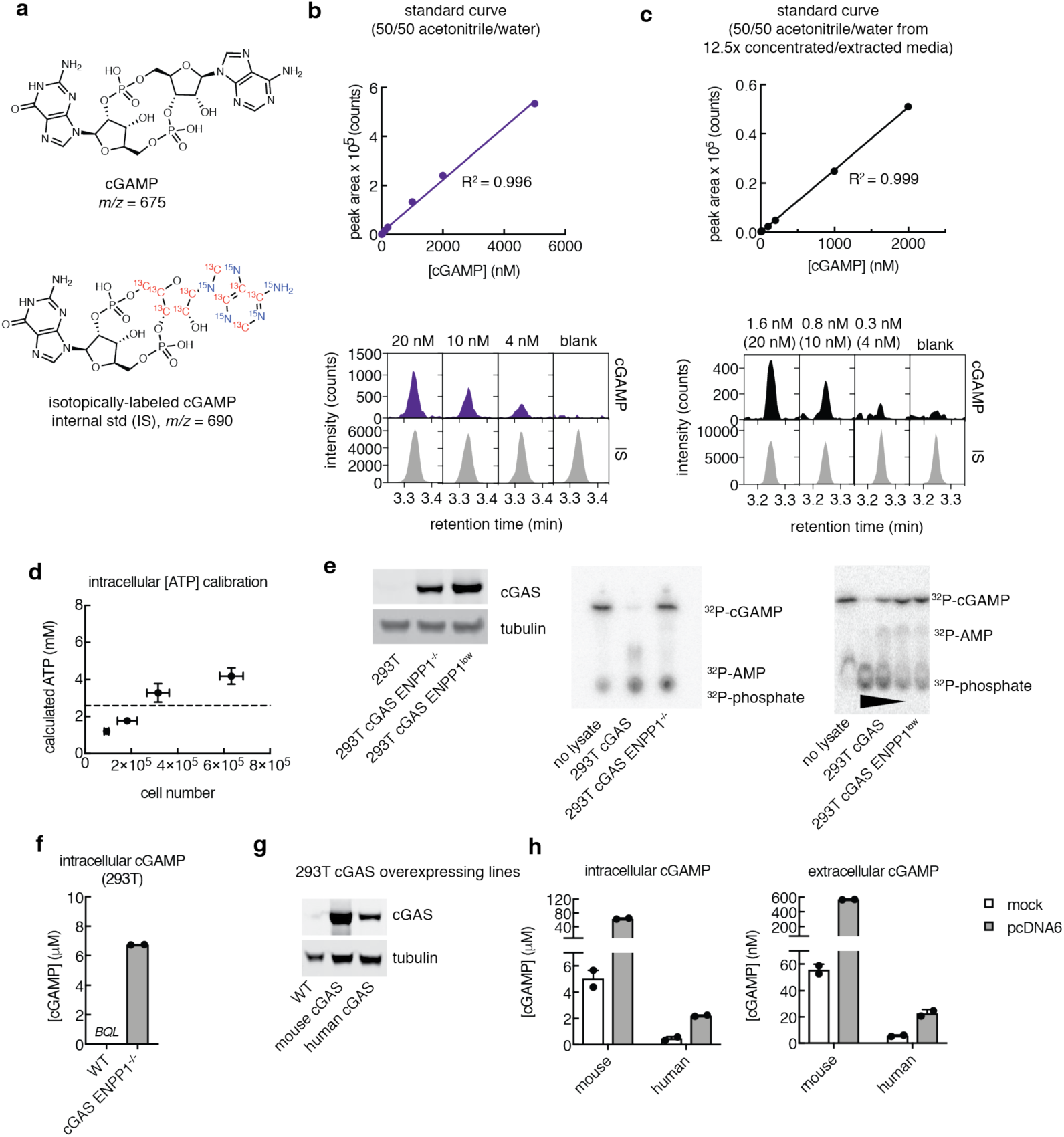
Measuring cGAMP in 293T cGAS ENPP1^low^ and 293T cGAS ENPP1^-/-^ cell lines by LC-MS/MS. **a**, Chemical structures of cGAMP (top) and single isotopically-labeled cGAMP (bottom) used as an internal standard at a concentration of 0.5-1 μM. **b-c**, Full (7-8 point) standard curves and LC traces of lowest cGAMP standards in 50/50 acetonitrile/water spiked in directly (LOQ = 4 nM) (**b**) or after concentrating and extracting 12.5x from complete cell culture media (LOQ = 0.3 nM from original sample, 4 nM in concentrated sample) (**c**). IS = internal standard. Data are representative of > 10 independent experiments. **d**, Calibration of cell number to ATP concentration measured by LC-MS/MS. Mean ± SEM (*n* = 2). **e**, cGAS expression of 293T, 293T cGAS ENPP1^-/-^, and 293T cGAS ENPP1^low^ cell lines analyzed by western blot (left). ENPP1 hydrolysis activity of ^32^P-cGAMP in whole cell lysates from 1 million each of 293T cGAS, 293T cGAS ENPP1^-/-^, and 293T cGAS ENPP1^low^ cells, measured by TLC and autoradiography (right). Lysate data are representative of two independent experiments. **f**, Intracellular concentrations of cGAMP from 293T WT and 293T cGAS ENPP1*^-/-^* cells without exogenous stimulation at steady state measured using LC-MS/MS. Mean ± SEM (*n* = 2). BQL = below quantification limit. **g**, Expression of cGAS in 293T WT, 293T stably expressing mouse cGAS, and 293T stably expressing human cGAS assessed by western blot. **h**, Intracellular and extracellular concentrations of cGAMP from 293T stably expressing mouse cGAS and 293T stably expressing human cGAS. Cells were mock transfected with only lipid or transfected with lipid + 0.5 µg/mL empty vector pcDNA6. After 24 hours, cells were refreshed with serum-free media and incubated for another 24 hours before analysis. Mean ± SEM (*n* = 2).

**Extended Data Figure 2.**
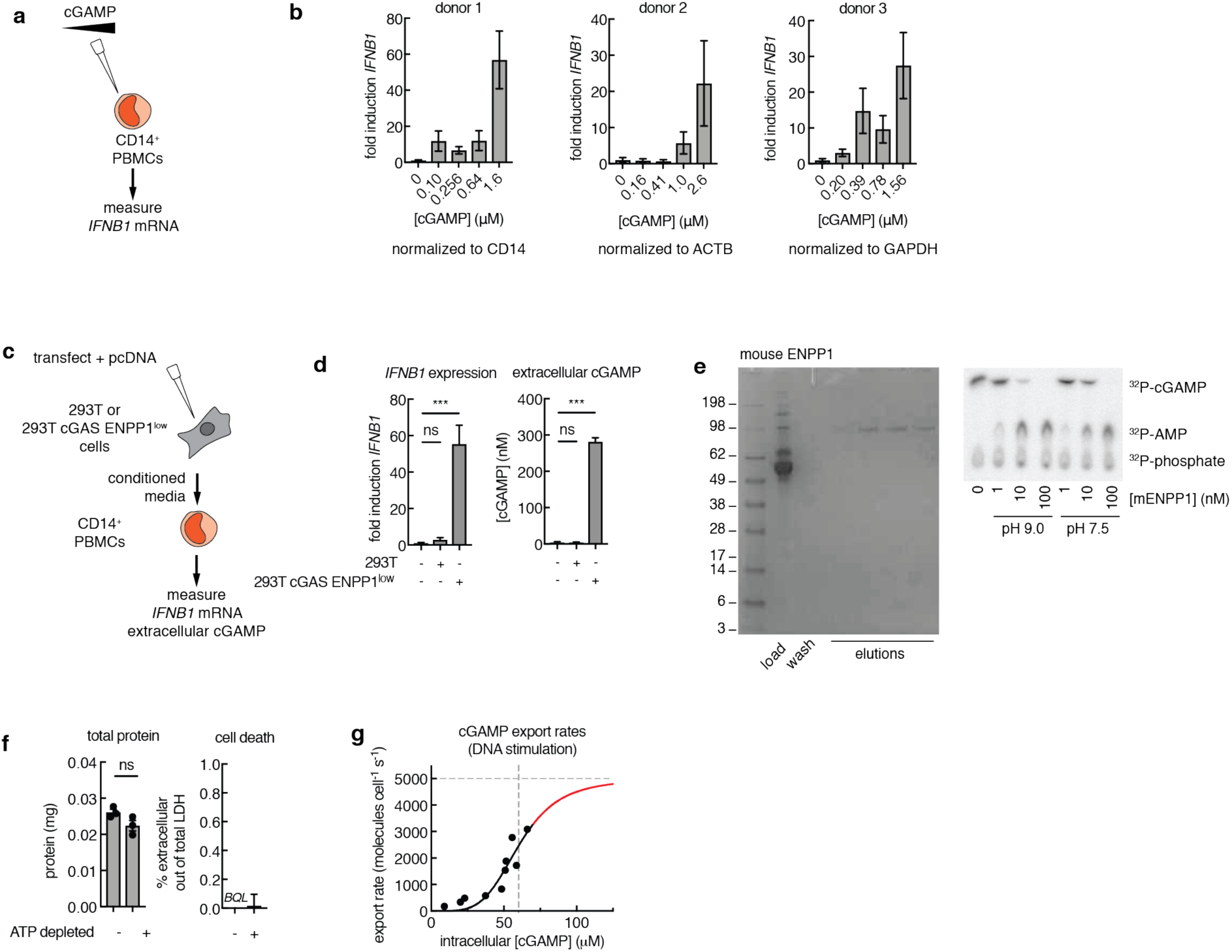
Characterization of cGAMP export by 293T cGAS ENPP1^-/-^ cells. **a**, Schematic of experiment for (**b**). Human CD14^+^ PMBCs stimulated with increasing concentrations of extracellular cGAMP for 16 h. **b**, *IFNB1* mRNA levels were normalized to indicated gene and fold induction was calculated relative to untreated CD14^+^ cells. Mean ± SEM (*n* = 2 technical qPCR replicates). **c**, Schematic of the conditioned media transfer experiment for (**d**). cGAS-null 293T or 293T cGAS ENPP1^low^ cells were transfected with 0.5 μg/mL empty pcDNA6 vector. Conditioned media from these cells was transferred to primary CD14^+^ human PBMCs and *IFNB1* expression was assessed after 16 h. **d**, *IFNB1* mRNA levels were normalized to *CD14* and the fold induction was calculated relative to untreated CD14^+^ cells. Mean ± SEM (*n* = 4). \*\*\**P* = 0.0003 (one-way ANOVA). cGAMP concentrations were measured in the conditioned media by LC-MS/MS. Mean ± SEM (*n* = 2). \*\*\**P* = 0.0002 (one-way ANOVA). Data are representative of two independent experiments. **e**, Coomassie gel of recombinant mouse ENPP1 purified from media; elution fractions were pooled before use (left). ^32^P-cGAMP degradation by mouse ENPP1 analyzed by TLC (right). **f**, 293T cGAS ENPP1^low^ cells were incubated with serum-free ATP depletion media (no glucose, 6 mM 2-deoxy-D-glucose, 5 mM NaN_3_) or serum-free complete media for 1 hour. Total protein content was measured by BCA and cell death was measured by lactate dehydrogenase activity. Mean ± SEM (*n* = 2-3). BQL = below quantitation limit. **g**, Data from Fig. 1l fit with the allosteric sigmoidal model *v_export_* = *V*_max_[*substrate*]*^n^* / (*K_half_*)*^n^* + [*substrate*]*^n^*, where *V*_max_ = the maximal transporter velocity, *n* = the Hill slope, and *K*_half_ = the substrate concentration at half *V*_max_. Here, we have constrained *V*_max_ < 5000 molecules cell^-1^ s^-1^, and the resulting fit shows that *K*_m_ > 60 μM.

**Extended Data Figure 3.**
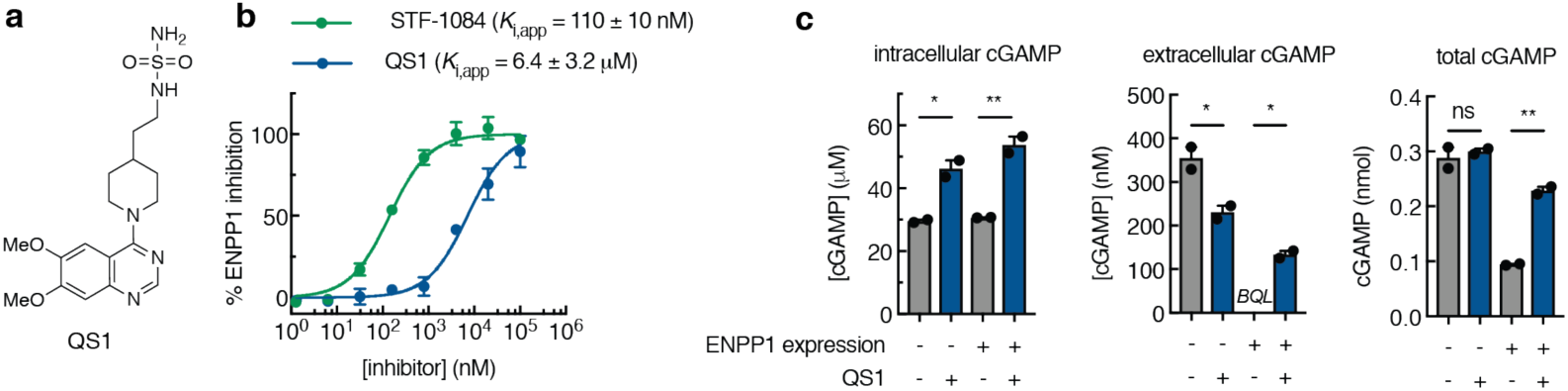
Improvement of STF-1084 over QS1. **a**, Structure of QS1. **b**, QS1 inhibitory activity (compared to STF-1084) against purified mouse ENPP1 with ^32^P-cGAMP as the substrate at pH 7.5 (QS1 *K*_i,app_ = 6.4 ± 3.2 μM). Mean ± SEM (*n* = 2 independent experiments). **c**, Intracellular, extracellular, and total cGAMP for 293T cGAS ENPP1*^-/-^* cells transfected with empty vector or vector containing human *ENPP1* in the presence or absence of QS1. cGAMP levels were measured after 24 hours by LC-MS/MS. Mean ± SEM (*n* = 2). \**P* < 0.05. \*\**P* < 0.01 (one-way ANOVA).

**Extended Data Figure 4.**
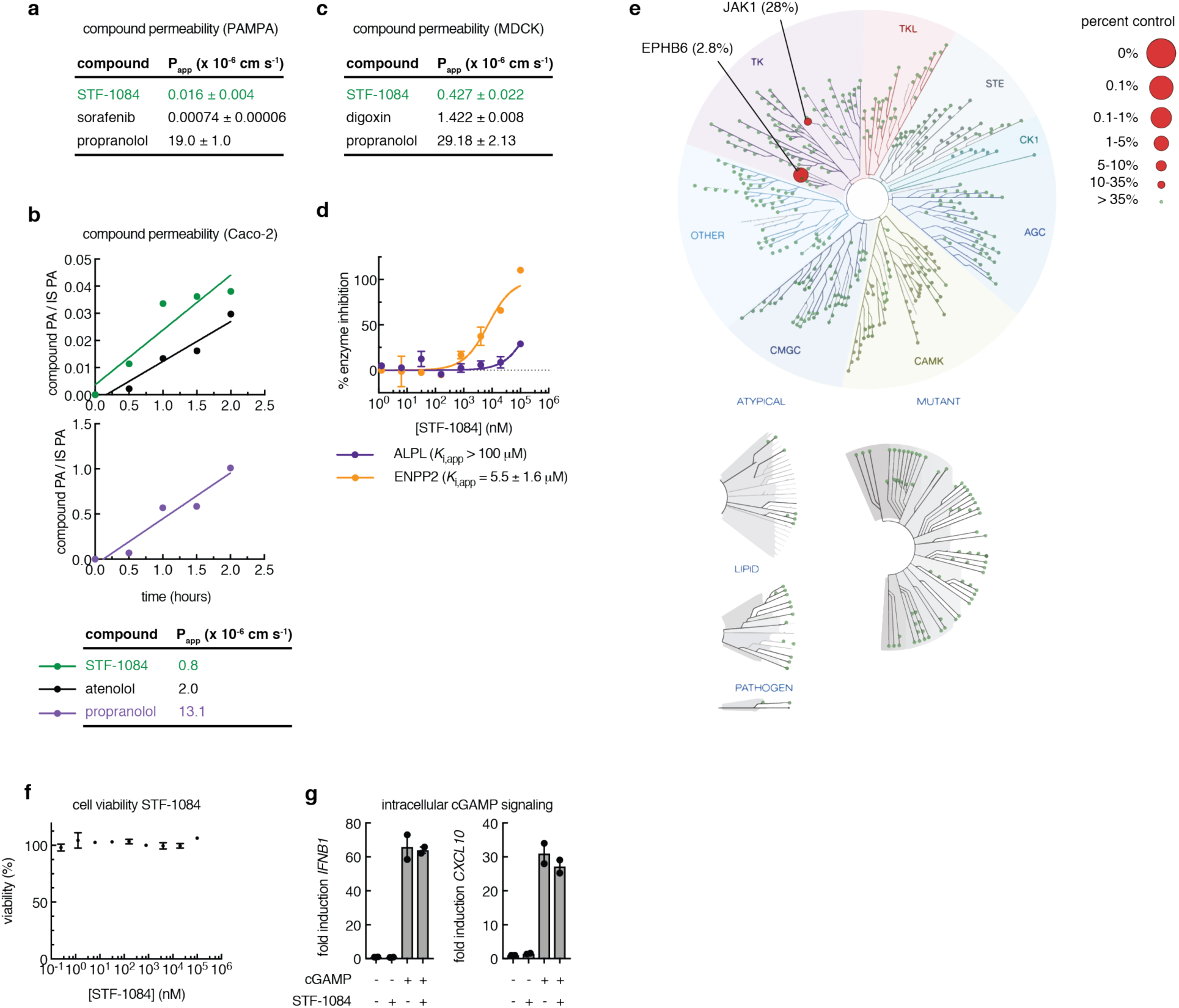
Development of the ENPP1 inhibitor STF-1084. **a**, Permeability of STF-1084 in artificial membrane permeability assay (PAMPA). **b**, Permeability of STF-1084 in intestinal cells Caco-2 assay. PA = peak area, IS = internal standard. Compounds, including STF-1084, atenolol (low passive permeability negative control) and propranolol (high passive permeability positive control), were incubated on the apical side of a Caco-2 monolayer for 2 hours. Compound concentration on the basolateral side was monitored by LC-MS/MS. Apparent permeability rates (P_app_) were calculated from the slope. Data are representative of two independent experiments. **c**, Permeability of STF-1084 in epithelial cells MDCK permeability assay. **d**, Inhibitory activity of STF-1084 against alkaline phosphatase (ALPL) and ENPP2. Mean ± SEM (*n* = 2). **e**, Kinome interaction map (468 kinases tested) for STF-1084 depicting kinase inhibition as a percent of control. **f**, Cell viability measured by CellTiterGlo. Total PBMCs were incubated with STF-1084 for 16 hours and then assayed for ATP levels using CellTiterGlo. Data was normalized to no STF-1084 to calculate % cell viability. Mean ± SEM (*n* = 2) with some error bars too small to visualize. **g**, Human PBMCs were mock electroporated or electroporated with 200 nM cGAMP and immediately incubated with media ± 50 μM of STF-1084 for 16 h. *IFNB1* and *CXCL10* mRNA levels were normalized to *ACTB* and the fold induction was calculated relative to untreated cells. Mean ± SEM (*n* = 2).

**Extended Data Figure 5.**
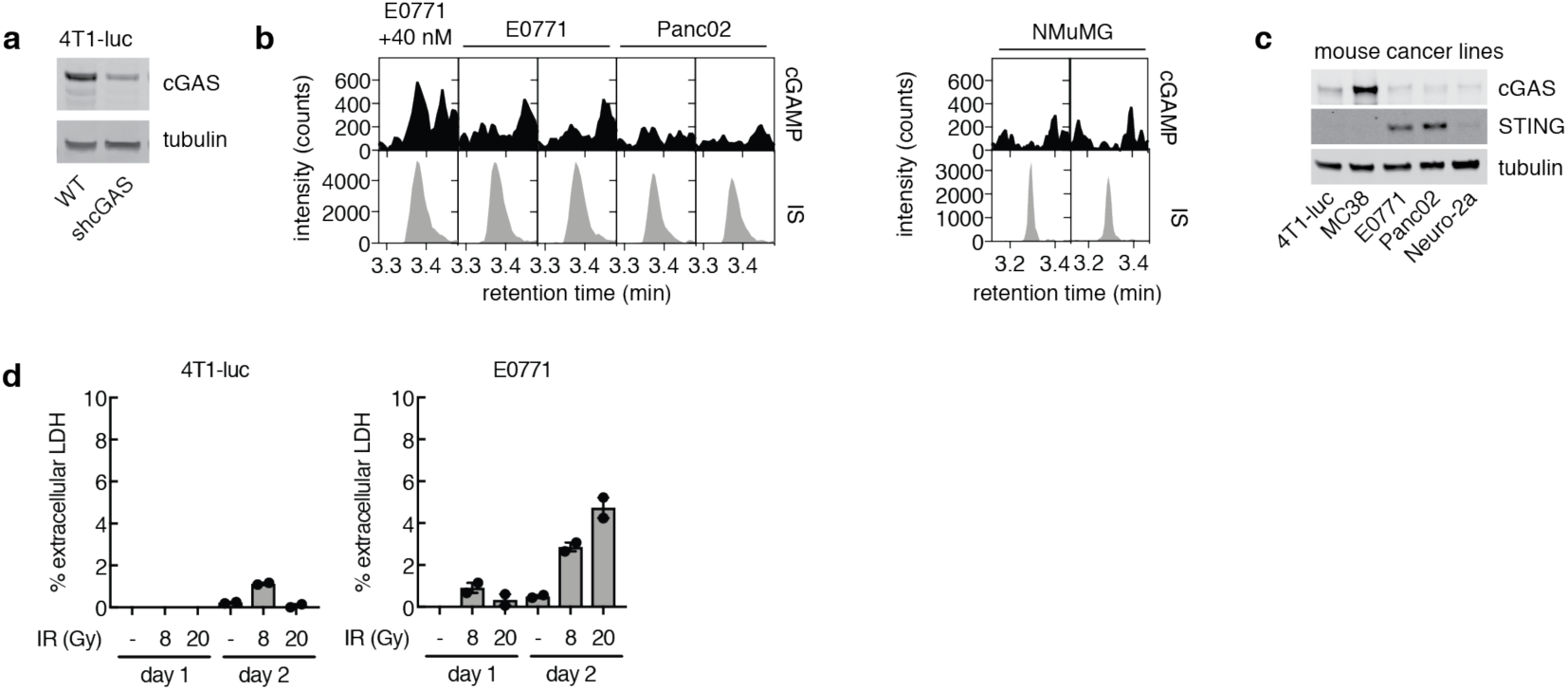
Cancer cell lines express cGAS and STING and remain viable with ionizing radiation (IR). **a**, cGAS expression of 4T1-luc WT and 4T1-luc shcGAS analyzed by western blot. **b**, Chromatograms for E0771, Panc02, and NMuMG cell lysate from LC-MS/MS. E0771 cell lysate was spiked with 40 nM cGAMP to determine limit of quantification. IS = internal standard. **c**, cGAS and STING expression of 4T1-luc, MC38, E0771, Panc02, and Neuro-2a cell lines analyzed by western blot. **b**, Cell viability of cancer cell lines 4T1-luc and E0771 measured by lactate dehydrogenase extracellular activity compared to intracellular activity. At time 0, cells were left untreated or treated with IR (8 Gy or 20 Gy) and refreshed with media supplemented with 50 μM STF-1084. Mean ± SEM (*n* = 2).

**Extended Data Figure 6.**
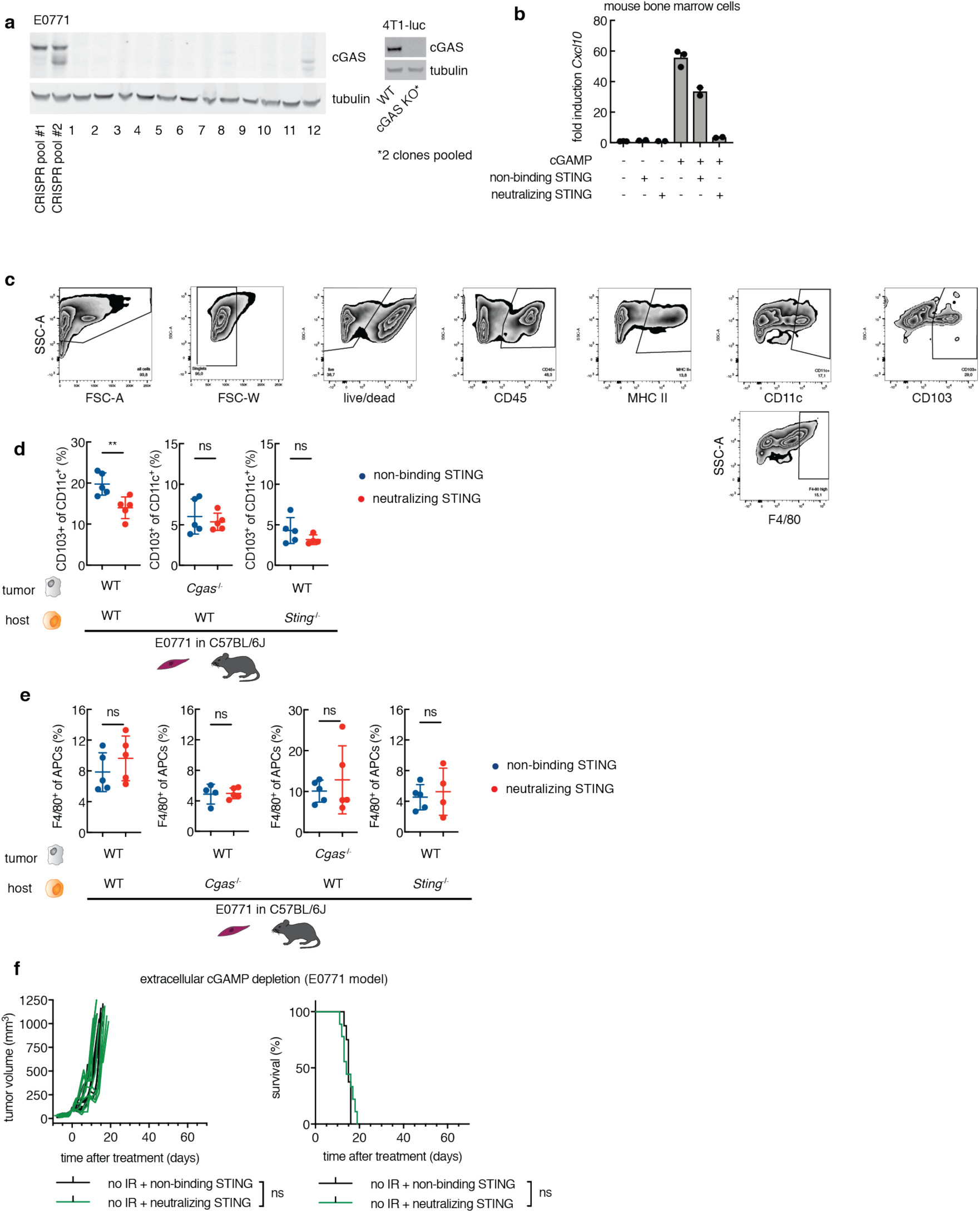
Sequestration of extracellular cGAMP decreases tumor-associated dendritic cells in a cancer cGAS and host STING dependent manner. **a**, E0771 (left) and 4T1-luc (right) *Cgas*^-/-^ cells subcloned from CRISPR knockout pools. Six E0771 *Cgas*^-/-^ subclones (1, 2, 4, 6, 8, and 9) were pooled before injection into mice. Two 4T1-luc *Cgas*^-/-^ subclones were pooled before injection into mice. **b**, *Cxcl10* mRNA fold induction (normalized to *Actb* and untreated cells) in primary mouse bone marrow cells treated with 20 μM cGAMP in the presence of neutralizing or non-binding STING (100 μM) for 16 h. Mean ± SEM (*n* = 2-3). **c**, FACS gating scheme for experiments in Fig. 4 f-h, Extended Data Fig. 6d, e. **d**, WT or *Cgas*^-/-^ E0771 cells (1×10^6^) were orthotopically injected into WT or *Sting*^-/-^ C57BL/6J mice on day 0. Neutralizing STING or non-binding STING was intratumorally injected on day 2. Tumors were harvested and analyzed by FACS on day 3. Samples were gated on cells in FSC-A/SSC-A, singlets (FSC-W), living cells, CD45^+^, MHC II^+^ (APCs), CD11c+ CD103+ (cDC1) population. (Non-binding STING: WT mice *n* = 5; *Cgas*^-/-^ cells *n* = 5; *Sting*^-/-^ mice *n* = 5. Neutralizing STING: WT mice *n* = 5; *Cgas*^-/-^ cells *n* = 5; *Sting*^-/-^ mice *n* = 4). Mean ± SD. \*\**P* < 0.01. **e**, WT or *Cgas*^-/-^ E0771 cells (1×10^6^) were orthotopically injected into WT, *Cgas*^-/-^ or *Sting*^-/-^ C57BL/6J mice on day 0. Neutralizing STING or non-binding STING was intratumorally injected on day 2. Tumors were harvested and analyzed by FACS on day 3. Samples were gated on cells in FSC-A/SSC-A, singlets (FSC-W), living cells, CD45^+^, MHC II^+^ (APCs), F4/80^+^ population. (Non-binding STING: WT mice *n* = 5; *Cgas*^-/-^ mice *n* = 4 *Cgas*^-/-^ cells *n* = 5; *Sting*^-/-^ mice *n* = 5. Neutralizing STING: WT mice *n* = 5; *Cgas*^-/-^ mice *n* = 5 *Cgas*^-/-^ cells *n* = 5; *Sting*^-/-^ mice *n* = 4). Mean ± SD. **f**, Established E0771 tumors (100 ± 20 mm^3^) were injected with non-binding (*n* = 8) or neutralizing STING (*n* = 9) every other day for the duration of the experiment. Tumor volume and survival were monitored.

**Extended Data Figure 7.**
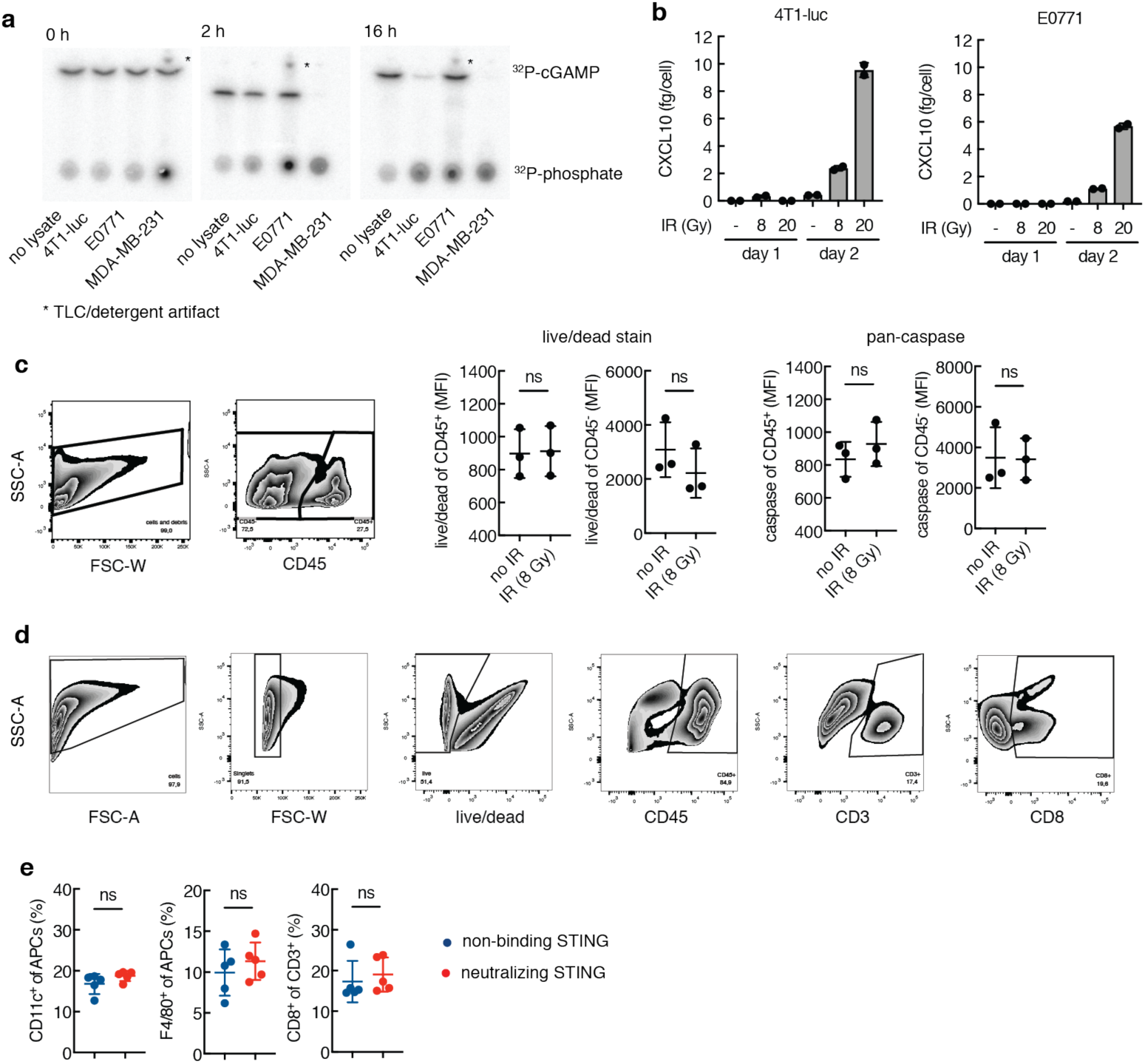
IR induces extracellular cGAMP-mediated CD8^+^ T cell activation without inducing cell death. **a**, ENPP1 activity in 4T1-luc, E0771, and MDA-MB231 cells using the ^32^P-cGAMP degradation assay. Data are representative of three independent experiments. **b**, CXCL10 production by cancer cell lines 4T1-luc and E0771 measured by ELISA. At time 0, cells were left untreated or treated with IR (8 Gy or 20 Gy) and refreshed with media supplemented with 50 μM STF-1084. Media was collected and cells were counted at indicated time points. Mean ± SEM (*n* = 2). **c**, FACS gating scheme for live dead analysis in established tumors. E0771 cells (5×10^4^) were orthotopically injected into WT C57BL/6J mice. The tumors were treated with IR (8 Gy) when they reached 100 ± 20 mm^3^ and harvested and analyzed by FACS 24h after IR. For caspase activity, a single cell suspension was incubated for 1h with the FAM-FLICA Poly Caspase substrate before FACS stain and analysis. Mean ± SD. **d**, FACS gating scheme for experiments in Fig. 4l, Extended Data Fig. 7e. **e**, E0771 cells (5×10^4^) were orthotopically injected into WT C57BL/6J mice. The tumors were treated with IR (8 Gy) when they reached 100 ± 20 mm^3^ and injected with non-binding (*n* = 5) or neutralizing STING (*n* = 5) on days 2 and 4 after IR. Tumors were harvested and analyzed by FACS on day 5. Mean ± SD.

**Extended Data Figure 8.**
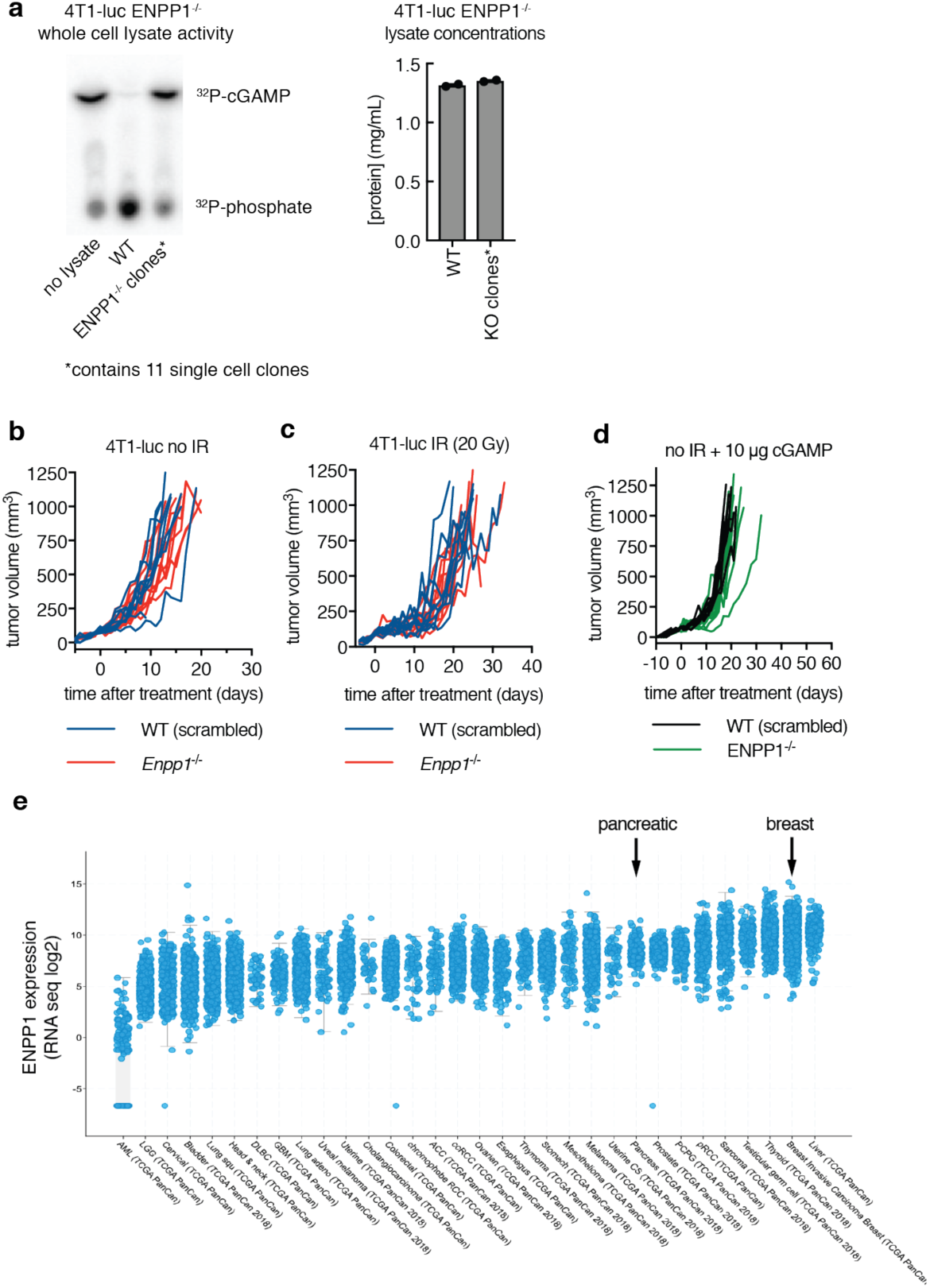
ENPP1 inhibition synergizes with IR treatment to increase tumor-associated dendritic cells. **a**, Validating *Enpp1*^-/-^ 4T1-luc clones (11 clones were pooled) using the ^32^P-cGAMP degradation assay. Lysates were normalized by protein concentrations. **b**, Established 4T1-luc WT (harboring scrambled sgRNA) (*n* = 10) or *Enpp1*^-/-^ tumors (*n* = 10) (100 ± 20 mm^3^) were monitored without treatment. Tumor volumes are shown. **c**, Established 4T1-luc WT (harboring scrambled sgRNA) (*n* = 10) or *Enpp1*^-/-^ tumors (*n* = 10) (100 ± 20 mm^3^) were treated with IR (20 Gy) and monitored. Tumor volumes are shown. **d**, Established 4T1-luc WT (harboring scrambled sgRNA) (*n* = 9) or *Enpp1*^-/-^ tumors (*n* = 9) (100 ± 20 mm^3^) were treated with three intratumoral injections of 10 µg cGAMP on day 2, 4, and 7 and monitored. Tumor volumes are shown. **e**, ENPP1 expression from RNA sequencing data from data generated by the TCGA Research Network: https://www.cancer.gov/tcga.

**Extended Data Figure 9.**
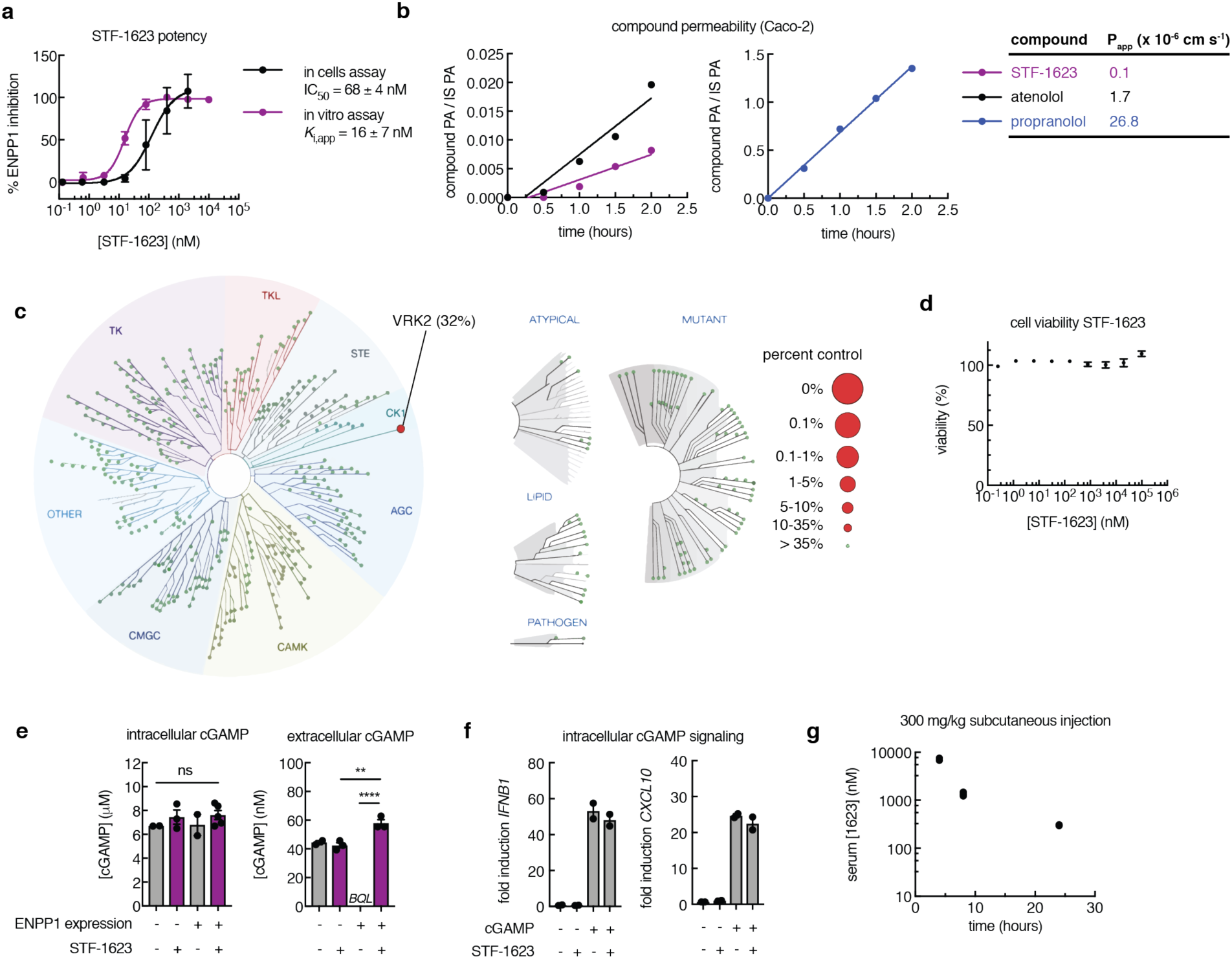
Development of the ENPP1 inhibitor STF-1623. **a**, Inhibitory activity of STF-1084 in vitro against purified mouse ENPP1 with ^32^P-cGAMP as the substrate at pH 7.5 (*K*_i,app_ = 110 ± 10 nM) and in cells against human ENPP1 transiently expressed in 293T cGAS ENPP1^-/-^ cells (IC_50_ = 340 ± 160 nM). Extracellular cGAMP levels were analyzed by LC-MS/MS after 24 hours. Mean ± SEM (*n* = 3 independent experiments for in vitro assay, *n* = 2 for in cells assay), with some error bars too small to visualize. **b**, Permeability of STF-1623 in intestinal cells Caco-2 assay. PA = peak area, IS = internal standard. Compounds, including STF-1623, atenolol (low passive permeability negative control) and propranolol (high passive permeability positive control), were incubated on the apical side of a Caco-2 monolayer for 2 hours. Compound concentration on the basolateral side was monitored by LC-MS/MS. Apparent permeability rates (P_app_) were calculated from the slope. Data are representative of two independent experiments. **c**, Kinome interaction map (468 kinases tested) for STF-1623 depicting kinase inhibition as a percent of control. **d**, Cell viability measured by CellTiterGlo. Total PBMCs were incubated with STF-1623 for 16 hours and then assayed for ATP levels using CellTiterGlo. Data was normalized to no STF-1623 to calculate % cell viability. Mean ± SEM (*n* = 2) with some error bars too small to visualize. **e**, Intracellular and extracellular cGAMP concentrations for 293T cGAS ENPP1^-/-^ cells transfected with empty pcDNA6 vector or vector containing human *ENPP1* in the presence or absence of 2 μM STF-1623 after 24 hours. cGAMP levels were analyzed by LC-MS/MS. *BQL* = below quantification limit. Mean ± SEM (*n* = 3). \*\**P* < 0.01. \*\*\*\**P* < 0.0001 (one-way ANOVA). Data are representative of two independent experiments. **f**, Human PBMCs were mock electroporated or electroporated with 200 nM cGAMP and immediately incubated with media ± 2 μM of STF-1623 for 16 h. *IFNB1* and *CXCL10* mRNA levels were normalized to *ACTB* and the fold induction was calculated relative to untreated cells. Mean ± SEM (*n* = 2). **g**, Mice were injected subcutaneously with 300 mg/kg STF-1623 at time 0. At indicated times, the mouse was sacrificed, blood was drawn by cardiac puncture, and serum isolated after clotting. STF-1623 concentrations were measure by LC-MS/MS. Individual data points shown (*n* = 2).

**Extended Data Figure 10.**
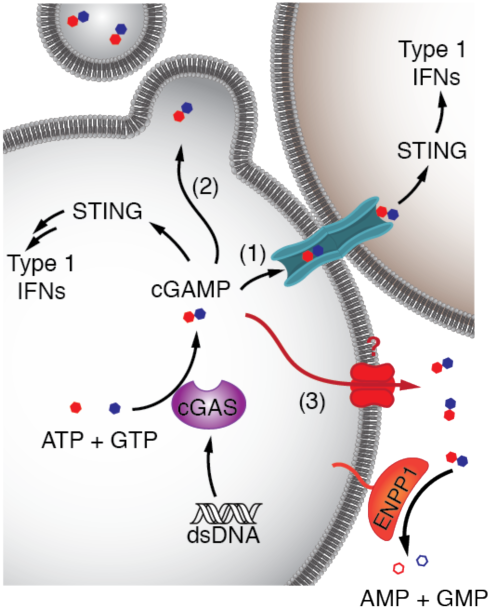
Different modes of cGAMP transmission from the synthesizing cell to target cells. (1) Spread via gap junctions; (2) packaged into budding viral particles and transmitted during the next round of infection; and (3) exported into the extracellular space.

**Extended Data Figure 11.**
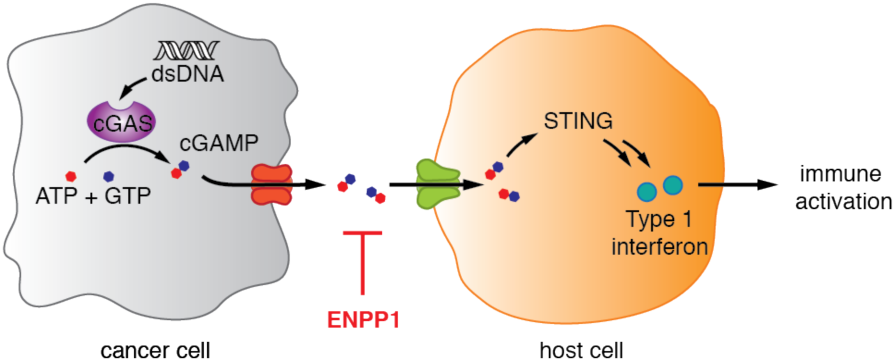
cGAMP is a cancer danger signal that can be harnessed by ENPP1 inhibition. Cancer cGAS detects its aberrant cytosolic dsDNA and synthesizes cGAMP. cGAMP is then exported into the extracellular space, where it can be degraded by the extracellular hydrolase ENPP1. Enhancing extracellular cGAMP by ionizing radiation or ENPP1 inhibition increases host detection and elimination of cancer cells.

